# Antibiotic Resistomes And Microbial Communities In The 18th-Century Urban Settlement And Slaughterhouse Environment

**DOI:** 10.64898/2026.07.31.742145

**Authors:** Minna Maria Maunula, Taru-Marja Mäkinen, Kari Uotila, Jenni Hultman, Kirill Bogdanov, Marko Virta, Johanna Muurinen

## Abstract

Antimicrobial resistance (AMR) is an ancient and natural phenomenon, yet it now poses a critical threat to global health. Human activities, particularly animal husbandry, have shaped microbial evolution by creating manure-rich environments that promote interactions between environmental and host-associated bacteria and facilitate horizontal gene transfer. Here, we investigated dormant, potential antibiotic-producing bacteria, microbial communities, and their AMR genes and mobile genetic elements in 18th-century preindustrial slaughterhouse surroundings excavated in Turku, Finland. By combining cultivation, genomic analyses, metagenomic sequencing, and ancient DNA authentication methods, we reconstructed preindustrial microbiomes and resistomes to better understand the early ecology and evolution of AMR, and to explore the role of antibiotic-producing bacteria in the emergence of AMR. Our results reveal putative ancestral forms of resistance mechanisms only recently characterized, such as fosfomycin thiol transferase *fosI,* plasmid-associated tmexCD-toprJ efflux pumps conferring resistance to last-resource antibiotic tigecycline, as well as sequences related to mobility of AMR genes. These findings demonstrate that key AMR elements were already present prior to widespread antibiotic use, reflecting their long-term environmental origins.

## Introduction

The ability of bacteria to resist antibiotics has become one of the greatest global public health and development threats, jeopardising the future of humanity. Although the widespread use of antibiotics since the early 1900s has significantly contributed to the emergence of AMR (Surette and Wright, 2017), antibiotic resistance is ancient and a natural phenomenon that arises from the co-evolution of antibiotic-producing bacteria and other bacteria in a shared environment (Berkner et al., 2014; D’Costa et al., 2011a). Over billions of years, bacteria have evolved to resist antibiotic molecules biosynthesized by antibiotic-producing microbes with antibiotic resistance genes (ARGs), and this has created a reservoir of resistance features in environmental bacteria, which is referred to as the resistome (Wright, 2010).

A major factor in facilitating the transfer of ARGs across different bacterial taxa and environments allowing bacteria to evolve to resist antibiotics, are mobile genetic elements (MGEs) (Gillings and Stokes, 2012a). Although transmission of ARGs from environmental resistome to pathogenic bacteria through MGEs is well established (Jiang et al., 2017; Peterson and Kaur, 2018a; Tamminen et al., 2012), in many cases, it is unclear how other environmental factors than selection pressure from the use of antibiotics contribute to it. Therefore, to understand the contemporary AMR crisis, it would be useful to analyze the co-existence of antibiotic producers and other bacteria in human-impacted environments before the antibiotic era.

Numerous clinically important antibiotics are originally products of environmental microorganisms, particularly actinomycetes such as *Streptomyces* (De Simeis and Serra, 2021). These spore-forming bacteria are common inhabitants of both natural and human-impacted environments, and antibiotic-producing *Streptomyces* strains have been isolated from diverse habitats, including soil, manure and flower beds (Schlatter and Kinkel, 2014). To avoid self-toxicity, antibiotic-producing bacteria often carry self-protecting genes that protect them from the compounds they synthesize (Mak et al., 2014). Many ARGs, such as genes for aminoglycoside targeting methylases and efflux pumps, have been shown to be transferred horizontally from antibiotic-producing bacteria to pathogenic bacteria even between phylogenetically distant bacteria, including transfers from Gram-positive to Gram-negative species (Courvalin, 2008). Besides horizontal gene transfer mediated by MGEs (Ghaly and Gillings, 2021), *Streptomyces* may share genetic material via mycelial networks (Berthold et al., 2016). While some ARGs carried by pathogens originate from antibiotic-producing microbes, others, such as CTX M β-lactamases and Qnr proteins, are believed to derive from non-producing environmental bacteria (Cantón, 2009). These transfer events are thought to be facilitated by environmental pollution following industrialization, which induced stress response mechanisms in environmental microbiomes and led to the enrichment of class 1 integron element (*intI1*). This element can capture and express genes that help bacteria survive under challenging conditions (Gaze et al., 2011; Gillings et al., 2015; Wright et al., 2008). First, *intI1* was identified in Enterobacteriaceae and other Gram-negative bacteria (Fluit and Schmitz, 1999) but later also reported in Gram-positive bacteria (Martin et al., 1990) and across a wide range of anthropogenically impacted environments (Gillings et al., 2015).

Since it is widely acknowledged that other human activities, such as intensive land use, pollution, and waste saturation have accelerated the spread of AMR (Gillings and Stokes, 2012a; Nolan et al., 2023), it would be important to clarify the role of environmental contamination before the antibiotic era. The emergence of agriculture has increased the prevalence of various types of manure and organic waste piles in human settlements. These piles also provide habitats for antibiotic-producing bacteria and with the constant input of MGE-carrying bacteria from humans and other animals as well as from the environment, organic waste piles may serve as gene-exchanging platforms, facilitating interactions that would not otherwise occur. Thus, such sites may have acted as hotspots for AMR propagation. Studying microbiomes and resistomes in these specific ecological niches, particularly in preindustrial settings could provide valuable insights into the evolution of AMR.

Over the recent years, microbial archeology combined with molecular methods such as metagenomic sequencing, together with recently developed ancient DNA (aDNA) based methods, have enabled us to shed light on the evolution of microbial communities and pathogenic bacteria (Kjær et al., 2022; Warinner et al., 2017; Wibowo et al., 2021a). These methods can also be applied to study the evolution of AMR from historical samples that contain both past microbiomes and ancient antibiotic-producing bacteria, which can be recovered as microbial seed banks capable of persisting in the environment for extremely long periods of time (Lennon and Jones, 2011). Since, archaeological samples typically contain a mixture of endogenous DNA from viable cells, which can be either metabolically active or dormant, and exogenous DNA from dead cells (Carini et al., 2016), it is essential to analyze both fractions to obtain a comprehensive view of microbiomes, particularly because soils generally harbor a mixture of aDNA and DNA in viable microbial cells (Carini et al., 2016).

Here, we investigated ARGs, MGEs, and microbial communities in archaeological samples originating from an 18th-century urban settlement and slaughterhouse environment uncovered during renovations of Turku Market Square, Finland, in winter 2022. These 18th-century layers were stratigraphically isolated from modern contamination, and well preserved due to sealing of by dense cobblestone paving, charcoal-rich deposits, and low-oxygen conditions that slowed decomposition. This setting provides a unique time window into an urban environment prior to the antibiotic era.

We collected a total of 12 samples representing human-impacted soils and various waste and fecal deposits. The study combines modern sequencing approaches with traditional cultivation methods, with a particular focus on capturing spore-forming, antibiotic-producing Gram-positive bacteria. We hypothesized that preindustrial human-impacted soils, and waste- and manure-rich environments served as hotspots for horizontal gene transfer, facilitating the integration of ARGs into MGEs through interactions among antibiotic-producing bacteria and bacteria associated with humans, other animals, and the environment. To address this hypothesis, we integrated cultivation-based and sequencing-based approaches to investigate the early emergence of AMR and to enlighten the role of antibiotic-producing bacteria in the ecology and evolution of AMR.

## Materials & Methods

### Study setting and sampling

Archaeological excavations were carried out during the renovation of Turku Market Square between 2018 and 2022 in Finland. The excavations uncovered parts of an old town, inhabited by craftsmen and merchants around the mid-17th century, demolished in the 1830s. Samples were collected in winter 2022 for both archaeological investigation and microbiological analyses, especially from a plot that differs greatly from others with a short but large-scale slaughterhouse activity in the turn of the 18th and 19th centuries. Samples contained soil before human impact, soils from the inhabited environment, and different kinds of wastes and manures, characterized by archaeologists participating in excavations. Samples for microbial analyses were collected aseptically in sterile plastic bags by microbiologists. For DNA extraction, samples were stored at -80°C in DNA extraction tubes and for cultivation at 6°C in double packed sterile plastic bags, as well as kept in dark and dry conditions. All sampled deposits, mostly dated to the 18th century, and their associated contextual information are summarized in Table 1.

**Table 1.** List of sampled deposits, and their associated contextual information.

| Sample ID | Origin | Matrix | Human Impact | Detailed information |
| --- | --- | --- | --- | --- |
| M9413_1 | Soil<br>(Timeserie 1) | Soil | Very low | Soil sample from a layer representing an older field or meadow phase before human activity emerged. |
| M9413_2 | Soil<br>(Timeserie 2) | Soil | Low | Soil sample from a layer representing an older field or meadow phase, located approximately 5 cm above the underlying sample. |
| M9413_3 | Soil<br>(Timeserie 3) | Soil | Low | Soil sample from a layer representing an older field or meadow phase, located approximately 5 cm above the underlying sample. |
| M9416 | Old field | Soil | High | Soil sample from a layer representing an older field or meadow phase before urbanization |
| M9409 | House yard | Soil | High | Soil sample collected from the yard area of a house next to the slaughterhouse area. |
| M9436 | Alley waste pit | Anthropogenic waste | High | Brown clay mixed with manure and wood waste, indicating waste disposal activity. |
| M9441 | Indoor spaces of the house | Anthropogenic waste | High | Material from inside the house, containing ash and possibly food-related remains. |
| M9438 | Garbage pit | Anthropogenic waste | High | Waste pit possibly for household waste. |
| M9419 | Barn waste pit | Anthropogenic waste | High | Clay mixed with wood waste and manure, indicating disposal of wastes related to livestock . |
| M9378 | Barn floor layer | Fecal | High | Manure-rich layer composed of straw and decomposed organic material, originating potentially from livestock. |
| M9352 | Defecation pit | Fecal | High | Manure-rich sediment from a defecation pit. |
| M9342 | Latrine | Fecal | High | Brown clay strongly mixed with fecal material, most likely from humans, since the sample was taken from a latrine. |

The archaeological remains were well preserved, given the typical soil conditions in Finland. Preservation of waste and manure pits was supported by sealing of the soil with dense paving of cobblestones in the 19th and 20th century. In addition, the sampled layers were also protected from microbial decomposition by charcoal rich layers from repeated fires, including the Great Fire of Turku in 1827. From redoximorphic features of the soil samples reflecting time before human impact, we determined that these layers had low oxygen conditions that slowed decomposition and the soil type (Visuri et al., 2021) further supported the preservation of DNA in these layers. Together, these features shielded the deposits from decomposition, water infiltration and other environmental disturbances. As a result, the 18th century layers remained stratigraphically isolated from modern contamination (Fig. 1). The 3D documentation of the excavation area is available as an interactive Matterport model https://my.matterport.com/show/?m=x9aku8BMUhB.

**Fig 1.**
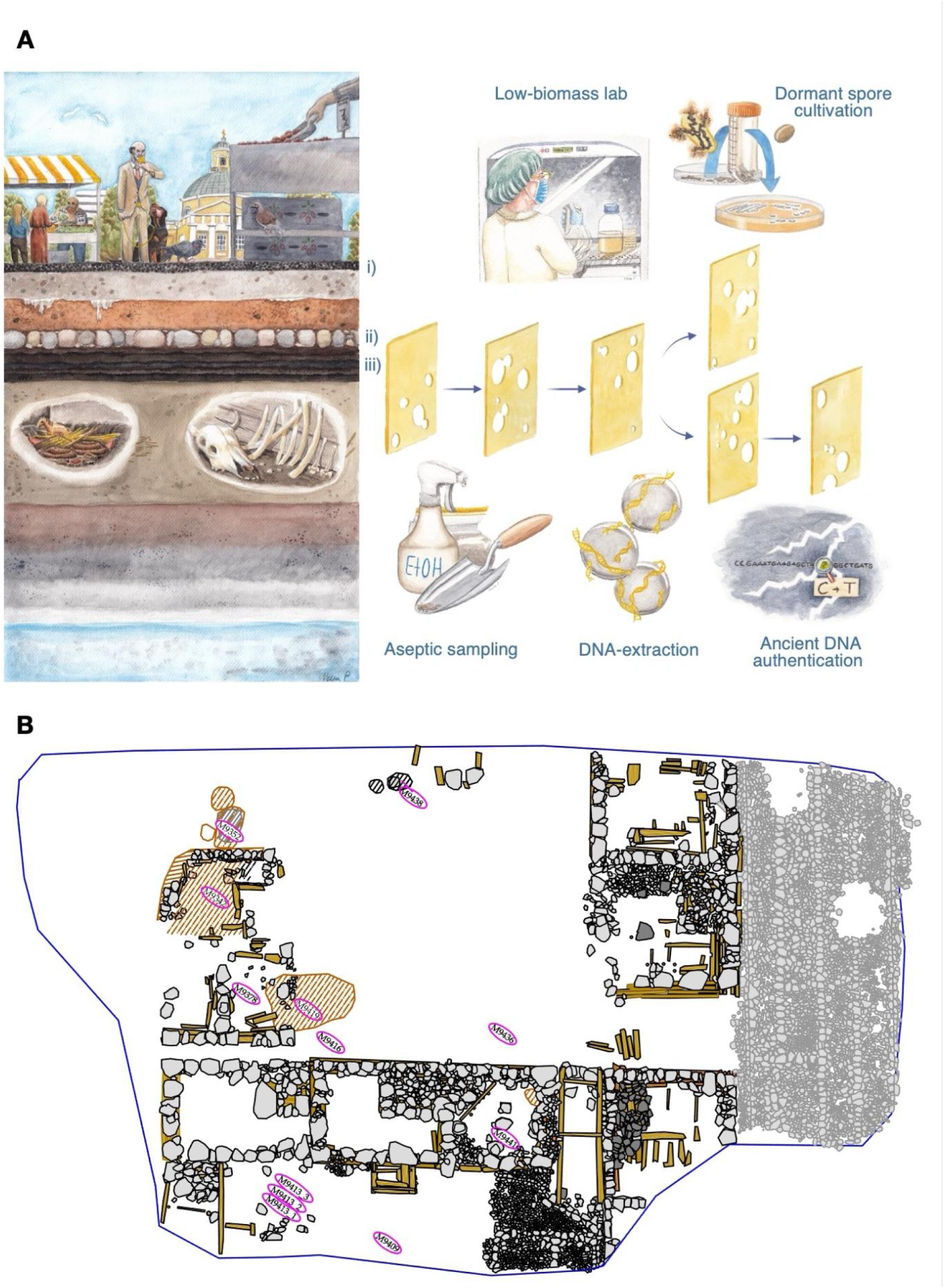
**(A)** Illustration of subsurface deposits beneath Turku Market Square showing the i) modern surface layer ii) historic cobblestones iii) a charcoal-rich burn layer from repeated urban fires, and beneath it an 18th-century cultural deposits associated with the slaughterhouse. These stratigraphically isolated, organic-rich layers contain waste pits, manure, and animal remains that support long-term preservation of microbial DNA, historic DNA and dormant spore-forming bacteria. **Next to this,** swiss-cheese model illustrating sequential physical, laboratory, and analytical barriers used to ensure authenticity of historical microbiomes and dormant spore-forming bacteria. **(B)** A sketch of the slaughterhouse environment, with the numbered sampling points circled in pink.

### Cultivation of Gram-positive bacteria

Inactivation of vegetative cells was carried out by drying approximately 15 ml of well-mixed sample material in a sterile petri dish and in a decontaminated laminar flow hood and for 24 hours. After drying, 3 g of the sample was loaded into a 50 ml conical centrifuge tube with 25 ml of sterile 1X phosphate buffered saline, vortexed for one minute at full speed and incubated for 30 minutes in a lateral shaking incubator at 50°C and 120 rpm to allow spore germination. Samples were then plated at a 10⁻⁴ dilution onto International Streptomyces project Medium No. 6 (ISP6) and7 (ISP7; without glycerol) (HiMedia Laboratories, Mumbai, India), and Gause’s No. 1 medium supplemented with a 1:1000 ratio of Nystatin (10,000 U/ml, Gibco, West Sussex, UK) to inhibit fungal growth and Nalidixic acid (30 μg/ml, Sigma-Aldrich, St. Louis, MO, USA) to suppress Gram-negative bacteria. Plates were incubated for approximately one week at 15°C and 28°C, and pure cultures were subsequently obtained. Pure cultures were stored in 20% glycerol stocks at −20°C until DNA extraction.

Bacterial isolates were re-cultivated from the glycerol stocks stored at −20°C. The isolates were grown at 28°C for approximately one week until a sufficient amount of biomass was obtained for DNA extraction.

### DNA extraction from environmental archaeological samples and isolated bacteria

Extractions of DNA were performed on archaeological soil samples using a modified hexadecyltrimethylammonium bromide (CTAB), phenol-chloroform, and bead-beating protocol (DeAngelis et al., 2009; Griffiths et al., 2000; Viitamäki et al., 2022) as follows: Approximately 0.5 g of sample material was added to 2 ml FastPrep Lysing Matrix E tubes (MP Biomedicals) and stored at −80°C until extraction. For each extraction, 0.5 ml CTAB extraction buffer was added, the tubes were vortexed briefly, and 0.5 ml phenol:chloroform:isoamyl alcohol (25:24:1) was added. Samples were vortexed laterally at full speed for 10 min, and centrifuged at 16,000 × g for 5 min at 4°C. The aqueous phase was transferred to 1.5 ml safe-lock Eppendorf tubes with 0.5 ml chloroform, vortexed, and centrifuged at 16,000 × g for 5 min at 4°C. The upper aqueous layer was transferred into tubes with 1 ml polyethylene glycol 6000 (PEG 6000) precipitation solution and incubated at room temperature for 1–2 h. A second extraction was performed by adding 0.5 ml CTAB buffer to the original Lysing Matrix E tubes and repeating all steps from phenol extraction through PEG precipitation.

Following precipitation, samples were centrifuged at 16,000 × g for 10 min at 4°C, and the supernatant was discarded. DNA pellets were washed 2–5 times with 0.5 ml cold 70% ethanol, each followed by centrifugation at 16,000 × g for 5 min at 4°C. Pellets were air-dried for ∼5 min, and resuspended in 25 µl C6 buffer (10mM Tris-HCl, pH 8.5; QIAGEN DNeasy® PowerLyzer® PowerSoil® Kit), after which the eluates from both extraction rounds were combined. A negative extraction control was included to monitor contamination.

Cleanup of extracted DNA was performed using AMPure XP magnetic beads (Beckman Coulter, Inc., Brea, CA, USA) to retain also aDNA fragments smaller than typically lost in column-based extractions. Beads were added to each sample at a 1.8× volume ratio, mixed thoroughly, and incubated for 10 min. Samples were then placed on a magnetic rack to separate the beads, washed twice with 0.5 ml 80% ethanol, and air-dried. DNA was eluted in 30 μl of C6 buffer (DNeasy® PowerLyzer® PowerSoil® Kit, QIAGEN, Hilden, Germany), gently vortexed, incubated for 5 min, and separated from the beads. Purified DNA was stored at −20°C until sequencing.

Genomic DNA from isolated bacteria was extracted using the DNeasy® PowerLyzer® PowerSoil® Kit (QIAGEN, Hilden, Germany) according to the manufacturer’s protocol, with the following modifications: the first incubation in step 7 was extended to 20 minutes, and the second incubation in step 9 was performed for 15 minutes. The extracted DNA was stored at −20°C until sequencing.

### Shotgun metagenomic & Whole-genome sequencing

Metagenomic sequencing for the DNA extracted from archaeological soil samples was carried out at the Institute of Biotechnology, University of Helsinki (Helsinki, Finland). Sequencing libraries were prepared by the institute’s sequencing facility and sequenced with the Aviti platform, producing 2 × 150 bp paired-end reads. To ensure sufficient sequencing depth, each sample was sequenced in two independent runs. Whole-genome sequencing of isolated bacterial strains was performed using the long-read PacBio Revio instrument (Pacific Biosciences, Menlo Park, CA, USA) with SMRTbell library preparation.

### Metagenomic data processing and analysis

#### Sequencing quality control and taxonomic classification

Raw sequencing reads were adapter-trimmed using FastP v0.24.0 (Chen et al., 2018), and reads shorter than 40 bp were discarded to ensure efficient sequence length for downstream analysis. Read quality was summarized using MultiQC v1.30 (Ewels et al., 2016). As each sample was sequenced across two runs, trimmed reads were concatenated per sample prior to downstream analyses. Taxonomic classification of metagenomic reads was performed with Kraken2 v2.1.5 (Wood et al., 2019) using a custom soil metagenomics database (db_vistamilk_soil_metagenomic_kraken2_2024; (Edwin et al., 2024). Relative abundances were re-estimated with Bracken v3.1 (Lu et al., 2017) at six taxonomic levels (phylum, class, order, family, genus, species) using a read length of 150 bp, a k-mer length of 35, and a minimum read assignment threshold of 10, meaning only taxa with at least 10 reads assigned by Kraken2 prior to re-estimation were included to filter out low-confidence detections.

#### Antibiotic resistance gene and mobile genetic element detection

Antibiotic resistance genes (ARGs) and mobile genetic elements (MGEs) were identified by searching trimmed reads against the ResFinder database v2.6.0 (Florensa et al., 2022) and a custom MGE database (Pärnänen et al., 2018), respectively, using BLASTN v2.17 (Altschul et al., 1990). Forward and reverse reads were combined per sample and converted from FASTQ to FASTA format prior to database searches. BLASTN was run with default word size (11) and an E-value threshold of 1×10⁻⁵.

#### aDNA authentication

Metagenomic assemblies required for downstream aDNA authentication were produced for each sample with MEGAHIT v1.2.9 (--min-contig-len 200) (Li et al., 2015). Three soil samples reflecting the time period before human activity (M9413_1, M9413_2, M9413_3) were additionally merged into a single combined sample for co-assembly, as individual per-sample mapping rates to their respective assemblies were low (30–40%). To assess the ancient origin of metagenomic sequences, contigs shorter than 1000 bp were first filtered out. Reads were then mapped to contigs using Bowtie2 v2.5.3 (--very-sensitive, -N 1) (Langmead and Salzberg, 2012). The resulting alignments were sorted using Samtools v1.21, and MD tags required for damage modelling were added to the sorted BAM files with samtools calmd, which encodes mismatches and deletions between the reference and read sequences in the alignment records (Danecek et al., 2021). DNA damage patterns were then estimated using pyDamage v1.0 with a damage modelling window of 30 bp (Borry et al., 2021). Contigs were considered ancient if they met all four criteria: q-value < 0.05, predicted 5’ damage (pmax) ≥ 0.01, minimum coverage ≥ 3×, and predicted accuracy ≥ 0.5. A pmax threshold of ≥ 0.01 was applied to account for the relatively recent depositional age of the samples (∼300–400 years back), in which DNA damage accumulation is expected to be limited compared to older archaeological specimens dating to thousands of years before present. Damaged and undamaged contigs were extracted separately from each assembly using seqtk.

To distinguish the authenticated aDNA from the whole metagenomic data, reads from each sample were mapped back to the damaged contigs of their respective sample using Bowtie2 v2.5.3 (--very-sensitive -N 1) (Langmead & Salzberg, 2012). Reads from three soil samples (M9413_1, M9413_2, M9413_3) were mapped to the damaged contigs against the co-assembly that was generated from these samples. Reads mapping to damaged contigs were extracted as paired FASTQ files using samtools fastq (-F 12) and subjected to the same taxonomic classification, ARG detection, and MGE detection pipelines as the whole-metagenome reads (described above). In addition, damaged contigs were annotated for ARGs using RGI v6.0.5 (-t contig -a DIAMOND --include_loose --local) against the canonical CARD database v4.0.1 (Alcock et al. 2023).

#### Statistical Analysis and Visualisation of the read based results

Statistical analysis and data visualizations were performed in R v4.5.3 using tidyverse package v2.0.0. Taxonomic community composition was visualized directly from Bracken relative abundance estimates, which re-estimate taxon abundances from Kraken2 classifications. Thus, both ARG and MGE read counts were normalized to Transcripts Per Million (TPM) by correcting for both gene length and sequencing depth by dividing raw counts by gene length in kilobases (reads per kilobase, RPK) and then scaling by the per-sample RPK sum to 10⁶. Reference gene lengths were extracted directly from the respective database FASTA files using the sequence lengths of all entries. BLASTN hits were retained at percentage identity ≥ 80%, alignment length ≥ 40 bp, and bitscore ≥ 50, retaining moderate- to high-confidence matches while removing weak, short, and low-quality alignments.

ARGs were annotated with resistance classes using the ResFinder phenotype table (phenotypes.txt), with multi-class entries consolidated into broader categories following target antibiotic drug class. Genes encoding efflux pump components (TOprJ, tmexD) were assigned to a separate Efflux/Multidrug class, and dihydrofolate reductase genes (dfr) and sulfonamide resistance genes (sul) were classified as Trimethoprim and Sulfonamide respectively. Prior to visualization and statistical analysis, ARG and MGE sequence variants were collapsed into canonical gene families by stripping trailing numeric variant suffixes, and TPM values were summed across variants within each family. MGE canonical gene families were classified into five categories: Integrase, Insertion Sequence, Transposase, Transposon-associated, and Plasmid-related.

Beta-diversity of ARG, MGE, and bacterial community profiles were analyzed with non-metric multidimensional scaling (NMDS, k = 2) on Hellinger-transformed Bray-Curtis dissimilarity matrices using the vegan package v2.7.1. For ordination and all downstream statistical analyses, the full gene family-level TPM matrices were used. Co-structure between resistome, microbiome, and mobilome ordinations were evaluated by symmetric Procrustes analysis and tested for significance using PROTEST with 999 permutations. Correlations between pairwise Bray-Curtis dissimilarity matrices were additionally assessed with Mantel tests using Pearson correlation and 999 permutations. Multiple testing correction was applied to both PROTEST and Mantel test p-values using the Benjamini-Hochberg procedure. All permutation-based analyses used a fixed random seed (set.seed(123)) for reproducibility.

### Whole genomic data processing and analysis

Adapter removal and filtering of raw PacBio HiFi reads were performed using HiFiAdapterFilt v2.0.1 (Sim et al., 2022). Read quality was assessed as for metagenomic data. Genome assemblies were generated with Hifiasm v0.19.5 (Cheng et al., 2021) using default parameters. Assembly quality statistics were calculated using QUAST v5.3.0 (Mikheenko et al., 2018). Genome completeness and contamination were assessed with CheckM2 v1.1.0 uniref100.KO diamond database (Chklovski et al., 2023). Assemblies showing signs of contamination or the presence of multiple species were binned using MetaBAT2 v2.15 (Kang et al., 2019). Taxonomic classification of assemblies and genome bins was performed with GTDB-Tk v2.4.0 (Chaumeil et al., 2022) with GTDB release 220 (Parks et al., 2020). Only genome bins and contigs of gram-positive isolates passing quality thresholds were retained for downstream analyses.

#### Metagenomic read mapping to isolate genomes

To estimate the relative abundance of the cultivated bacterial isolates within the metagenome, trimmed metagenomic reads were mapped against a combined reference of all isolate genome assemblies using Bowtie2 v2.5.3 (-D 20 -R 3 -N 1 -L 20 -i S,1,0.50) (Langmead and Salzberg, 2012). Per-isolate read counts were summarised into a count matrix using samtools v.1.21 idxstats (Danecek et al., 2021).

#### Antibiotic resistance gene and mobile genetic element detection

Antibiotic resistance genes were identified by aligning assembled genomes or bins against the ResFinder database v2.1.1 (Florensa et al., 2022) using BLASTN v2.17.0 (-evalue 1e-5 -perc_identity 70 -word_size 11). Mobile genetic elements were identified by aligning assembled genomes or bins against the custom MGE database (Pärnänen et al., 2018) using BLASTN v2.17.0 with the same parameters as for ARGs.

#### Antibiotic related biosynthetic gene cluster detection

The antibiotic production potential of the isolates was assessed by identifying secondary metabolite biosynthetic gene clusters (BGCs) using AntiSMASH v8.0.4 (Blin et al., 2025). Prodigal was used as a gene prediction tool (--genefinding-tool prodigal) and cluster comparisons were performed against the Minimum Information about a Biosynthetic Gene Cluster database (MIBiG) database v4.0 (--cc-mibig) (Zdouc et al., 2025). AntiSMASH output was parsed using a custom Python script to extract BGC region annotations and their closest MIBiG matches. BGC results were further analyzed in R using tidyverse v2.0.0 and jsonlite v2.0.0 packages. BGCs were classified as antibiotic-producing if their most similar known cluster was annotated with antibacterial activity, as defined by MIBiG bioactivity terms. Only BGCs hits with ≥15% similarity to a reference cluster annotated with antibacterial activity in MIBiG were retained for visualisation.

#### Visualisation of the isolate results

BLASTN output files for ARGs and MGEs were parsed in R v4.5.3. using dplyr v1.1.4. and ggplot2 v3.5.2. packages. For each isolate, overlapping hits on the same contig were resolved by retaining the hit with the highest percentage identity. Gene coverage was calculated as the alignment length divided by the full reference gene length. For ARGs, hits were retained if they met thresholds of ≥70% nucleotide identity and ≥60% gene coverage; for MGEs, thresholds of ≥70% identity and ≥40% coverage were applied. ARGs were manually annotated by antibiotic class based on gene name prefixes. Antibiotic-production-associated BGCs were visualized with as heatmaps showing percentages per isolate, produced with ggplot2.

## Results

### Microbial community composition differs between the entire and the damaged metagenomic dataset in the environment of an 18th-century slaughterhouse

We obtained in total 1563 million raw reads, and in total of 1551.2 million reads after adapter removal and quality trimming from the 12 studies samples. Between 19-49% of the sample reads were taxonomically classified against a custom-built microbial kraken2 database by Edwin et al. 2024 (Supplementary Table 1). Bacterial sequences comprised 94.85% of the metagenomic reads. Archaeal sequences constituted 1.80% of the total reads, with Euryarchaeota (1.79%) being the most abundant archaeal taxa. Eukaryotic sequences accounted for 3.35% of reads, all of which were fungal in origin, dominated by Ascomycota (2.41%) and Basidiomycota (0.94%).

Across all metagenomic sequences, the most abundant phyla were Pseudomonadota (40.8%), Actinomycetota (12.6%), Bacillota (7.16%), Bacteroidota (6.39%), and Chloroflexota (3.40%). The dominant orders were Pseudomonadales (15.8 %), Burkholderiales (8.28%), Moraxellales (4.92%), Hyphomicrobiales (3.51%), and Flavobacteriales (3.43%), although their relative abundances varied among samples (Fig. 2A). All the samples shared similar patterns in order level, except sample M9441 from indoor spaces of a house containing ash, with a higher proportion of Pseudomonadales and Burkholderiales. The most abundant genera differed by environment type (Fig. 3A); however, overall, the major genera across all sequences were *Pseudomonas* 15.9%, *Acinetobacter* 2.88%, *Flavobacterium* 2.83%, *Janthinobacterium* 2.32%, and *Candidatus Aegiribacteria* 2.27%.

**Fig 2.**
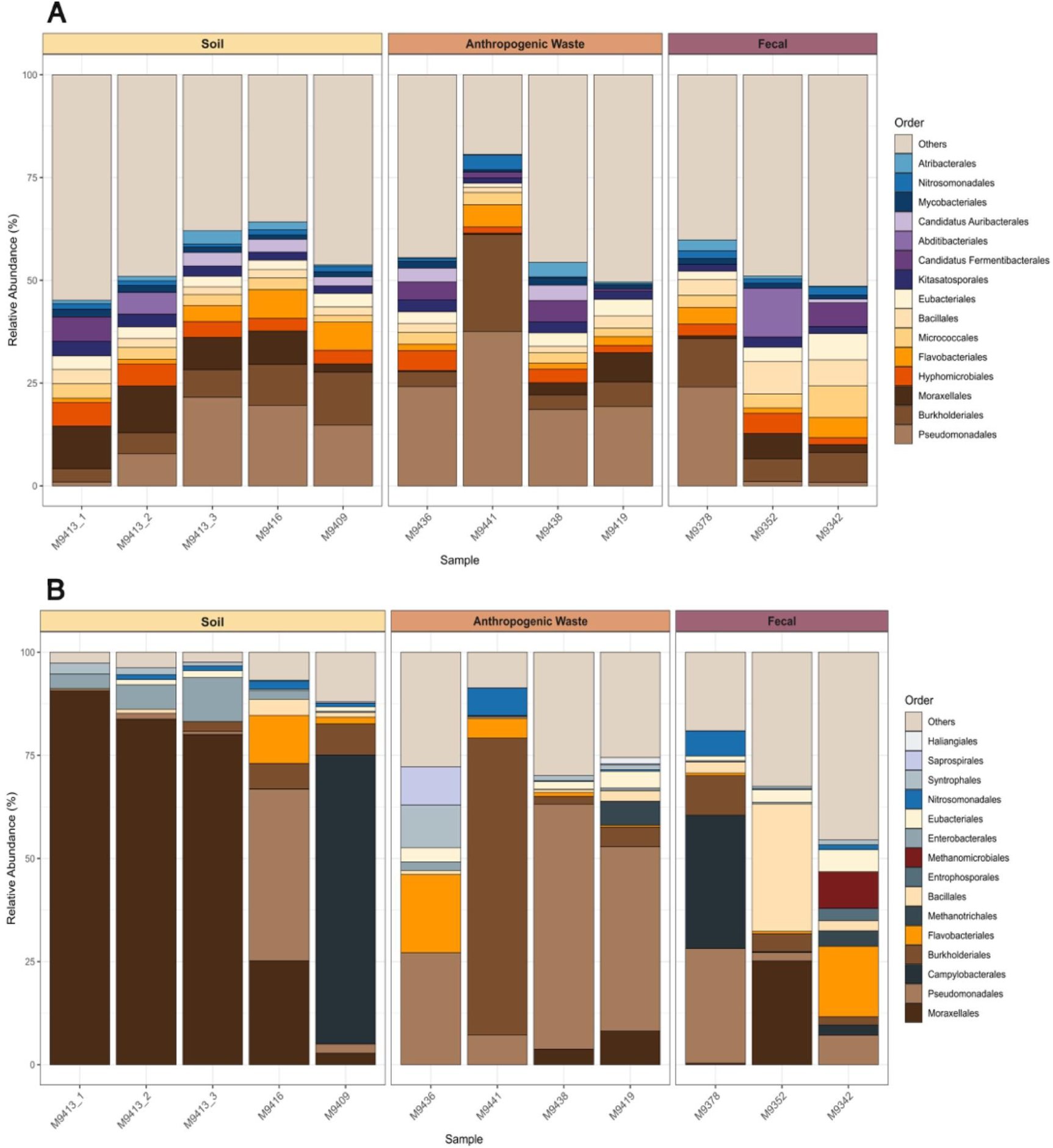
Taxonomic composition of reads classified with Kraken2 and normalized with Bracken at the order level, showing the relative abundance of the 15 most abundant microbial orders across metagenomes of 12 archaeological samples. The data are shown for (A) the complete metagenomic dataset and (B) ancient metagenomic reads mapped to pyDamage-authenticated damaged contigs. Samples are displayed along the x-axis, and relative abundance estimates as percentages based on Bracken-corrected read counts are shown on the y-axis. Orders outside the top 15 most abundant were grouped into an “Others” category. Samples are grouped by the main contents of the archaeological sample material (Soil, Anthropogenic Waste, and Fecal). Each microbial order is represented by a unique color; orders present in both datasets are shown using the same color in both panels.

**Fig 3.**
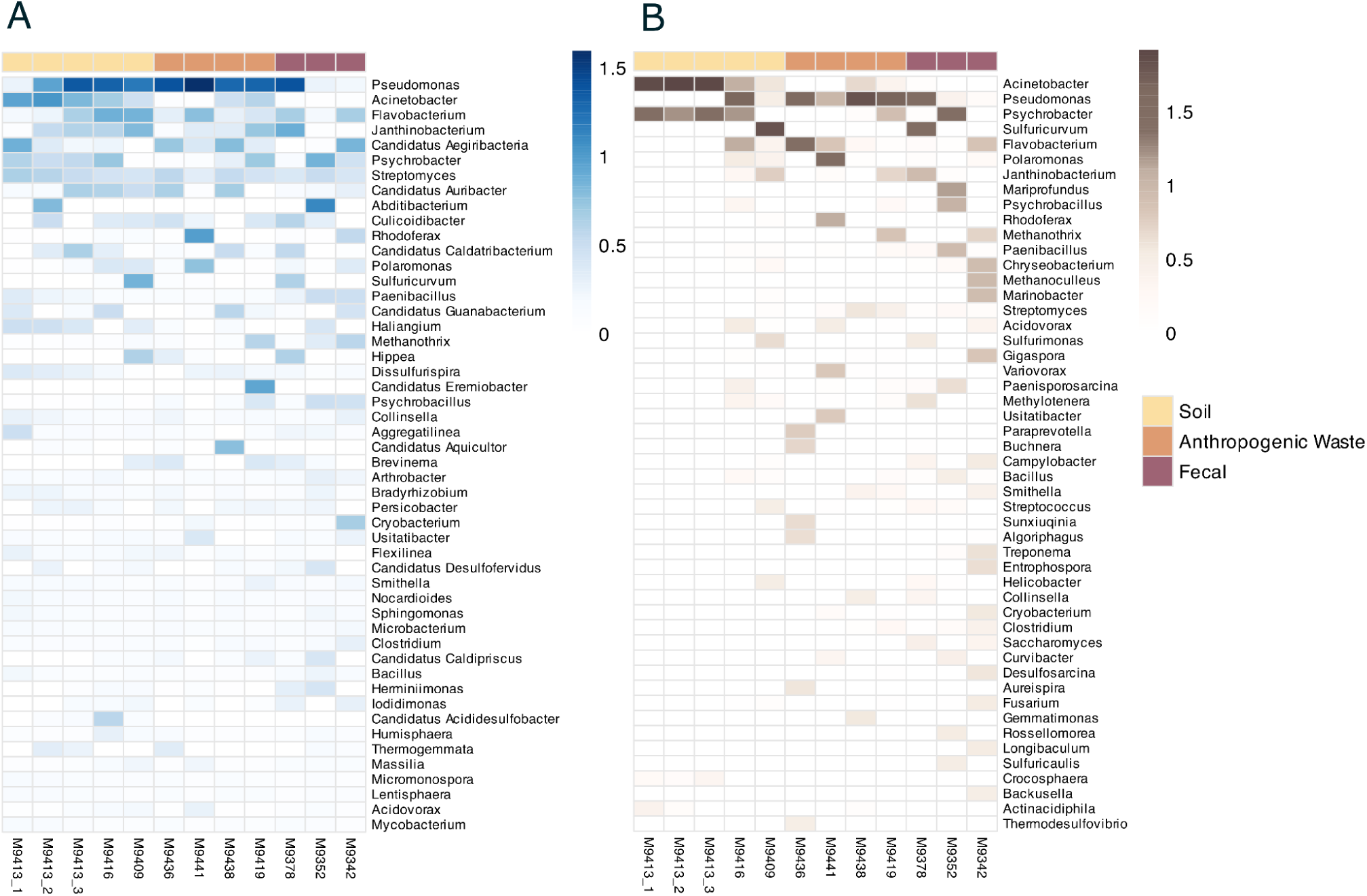
Heatmap showing the top 50 most abundant microbial genera across 12 archaeological samples in (A) the complete metagenomic dataset and (B) ancient metagenomic reads mapped to pyDamage-authenticated damaged contigs. The three colors at the top of the heatmap indicate the main contents of the archaeological sample material. Samples are displayed on the x-axis and detected genera on the y-axis. Taxonomic classification for reads was performed with Kraken2 and normalized using Bracken at the genus level. The heatmap scale represents log_10_-transformed relative abundance (log_10_(% + 1)). Genera are ordered by overall abundance, with the most abundant displayed at the top. White cells indicate genera not detected in a sample.

We extracted a total of 2200 damaged contigs using pyDamage, and after mapping metagenomic reads into the damaged contigs, we obtained a total of 1,973,394 damaged reads. Of these, 96.98% were identified as bacterial in origin. Archaeal sequences represented 1.30% of the ancient read dataset and were exclusively assigned to Euryarchaeota, with Methanomicrobia accounting for 1.17% of the total reads. Eukaryotic sequences comprised 1.72% of the ancient reads, all of which belonged to fungal taxa, predominantly Ascomycota (1.36%) and Basidiomycota (0.36%).

Across all ancient reads, the most abundant phyla were Pseudomonadota (62.2%), Campylobacterota (16.2%), Bacillota (4.51%), Bacteroidota (3.62%) and Actinomycetota (3.61%). At the order level Burkholderiales (29.7%), Pseudomonadales (20.7%), Campylobacterales (16.7%) dominated the dataset, followed by Nitrosomonadales (4.37%) and Moraxellales (3.63%). The overall variation in order levels was more notable in the damaged dataset (Fig. 2B). At the sample level, samples M9413_1-3, reflecting the soil prior to urbanized anthropogenic activity, exhibited a relative abundance of over 75% for Moraxellales (from phylum Pseudomonadota), making it the most dominant order. Interestingly, sample M9409, collected from a slaughterhouse yard, showed over 70% abundance of Campylobacterales (phylum Proteobacteria, Fig. 2B), whereas this order was not particularly prominent in the overall metagenomic data (Fig. 2A). Additionally, Campylobacterales were detected at approximately 25% relative abundance in sample M9378, which originated from the barn floor. Burkholderiales were notably prominent, with over 70% relative abundance in damaged reads from M9441, but also over 20% of the entire metagenomic data. The dominant genera in the ancient read dataset were annotated as Pseudomonas (23.2%), Sulfuricurvum (15.7%), Polaromonas (9.4%), Janthinobacterium (3.85%) and Rhodoferax (3.56%) (Fig. 3B). The ancient reads contained a greater diversity of bacterial genera, including genera with clinically relevant pathogens such as *Treponema, Campylobacter, Streptococcus*, and *Helicobacter.* However, these pathogens were primarily observed in individual samples and were not consistently shared across the dataset.

After the metagenomic assembly, a total of 2,739,045 contigs were identified. As stated above, we detected a total of 2,200 damaged contigs, of which 1,682 contigs had damage levels ranging from 1% to 5%, 378 contigs with damage levels between 5% and 10%, and 140 contigs exhibiting damage levels exceeding 10% (Supplementary Fig. 1). Sample M9419 collected from the waste pit inside the barn exhibited the highest number of damaged contigs (n=630), followed by samples M9378 (manure from the barn), M9441 (sample containing ash taken from inside the house), and M942 (fecal material from the latrine). The total number of contigs was not correlated with the number of damaged contigs (Spearman’s rho = 0.236, p = 5.14e-01). After extracting the damaged contigs, only 0.04-2.01% of the metagenomic reads mapped back to these contigs, depending on the sample (Supplementary Fig. 1 and Supplementary Table 2).

### Anthropogenic influence and waste disposal leave traces in antibiotic resistance gene and mobile genetic element profiles

A total of 1,741,338 BLASTN hits were detected after mapping metagenomic reads against the ResFinder database (Ferrer Florensa et al. 2022). After quality filtering (percent identity ≥ 80%, alignment length ≥ 40 bp, bitscore ≥ 50), 530,808 hits were retained. Following the collapsing of sequence variants into canonical gene families, 543 unique ARG families were identified that were falling into 16 distinct antibiotic resistance classes. Beta-lactams represented the largest proportion of detected ARG families (24.68%), followed by aminoglycosides (24.31%), macrolide-lincosamide-streptogramin (MLS) (15.65%), tetracyclines (9.94%), phenicols (6.45%), fosfomycin (4.05%), trimethoprim (4.05%), glycopeptides (2.58%), polymyxins (1.84%), quinolones (1.66%), efflux/multidrug (2.21%), other (1.10%), sulfonamides (0.55%), fusidic acid (0.37%), mupirocin (0.37%), and rifamycins (0.18%). The ten most abundant ARG families across all metagenomic samples were *blaOXA, ole(C), tmexD2, tmexD4, tmexD3, tmexD1, blaSHV, ole(B), tlr(C)*, and *OqxB*.

Overview of the top 50 most abundant ARG families revealed only mild differences in distribution patterns across sample types (Fig. 4A). Aminoglycoside resistance genes *aadA11, aadA16*, and *aadA6* were consistently detected at high abundance across all samples. Beta-lactamase genes showed variability between samples, with several absent from the low human impact soil sample M9413_1. Among the beta-lactam class, *bla*_CARB_ and *bla*_TEM_ were notably less abundant than other *bla* family genes, and *bla*_TEM_ was detected only in M9413_1 and M9438 originating from the garbage pit. Efflux/multidrug resistance genes *TOprJ1–4*, *tmexD1–4*, and *tmexC1–4*, encoding the TmexCD-TOprJ Resistance-Nodulation-Cell Division (RND)-type efflux pump system, were detected across all samples. Of these, *tmexD* variants were the most abundant, followed by *TOprJ*, while *tmexC* variants were consistently present at lower abundance and were not included among the 50 most abundant ARG families shown in the heatmap (Fig. 4A). Fosfomycin resistance genes *fosA* and *fosA3*–*7*, which can be chromosomally encoded or carried on plasmids via IS*26*, were detected in multiple samples, with *fosA* present in all samples while *fosA3*–*7* were absent from M9352, M9342, and M9413_1. Six MLS resistance gene families were identified, of which *mph*(E) was detected in five out of 12 samples, while the remaining five were present in all samples. The sulfonamide resistance gene *sul4* was detected in every sample, whereas the trimethoprim resistance gene *dfrA26* was absent from M9413_1 and M9342. The plasmid-mediated quinolone resistance gene *qepA4*, encoding a QepA4 efflux pump, was detected across samples and likely reflects an environmental ancestor gene that has since evolved into a clinically relevant ARG. The mupirocin resistance gene *mupA* was present in all samples. Tetracycline resistance genes *otr*(A), *otr*(C), *tcr3*, *tet*, and *tet*(Q) were detected across all metagenomic samples, with the single exception of *tet*(Q), which was absent from M9413_1.

**Fig 4.**
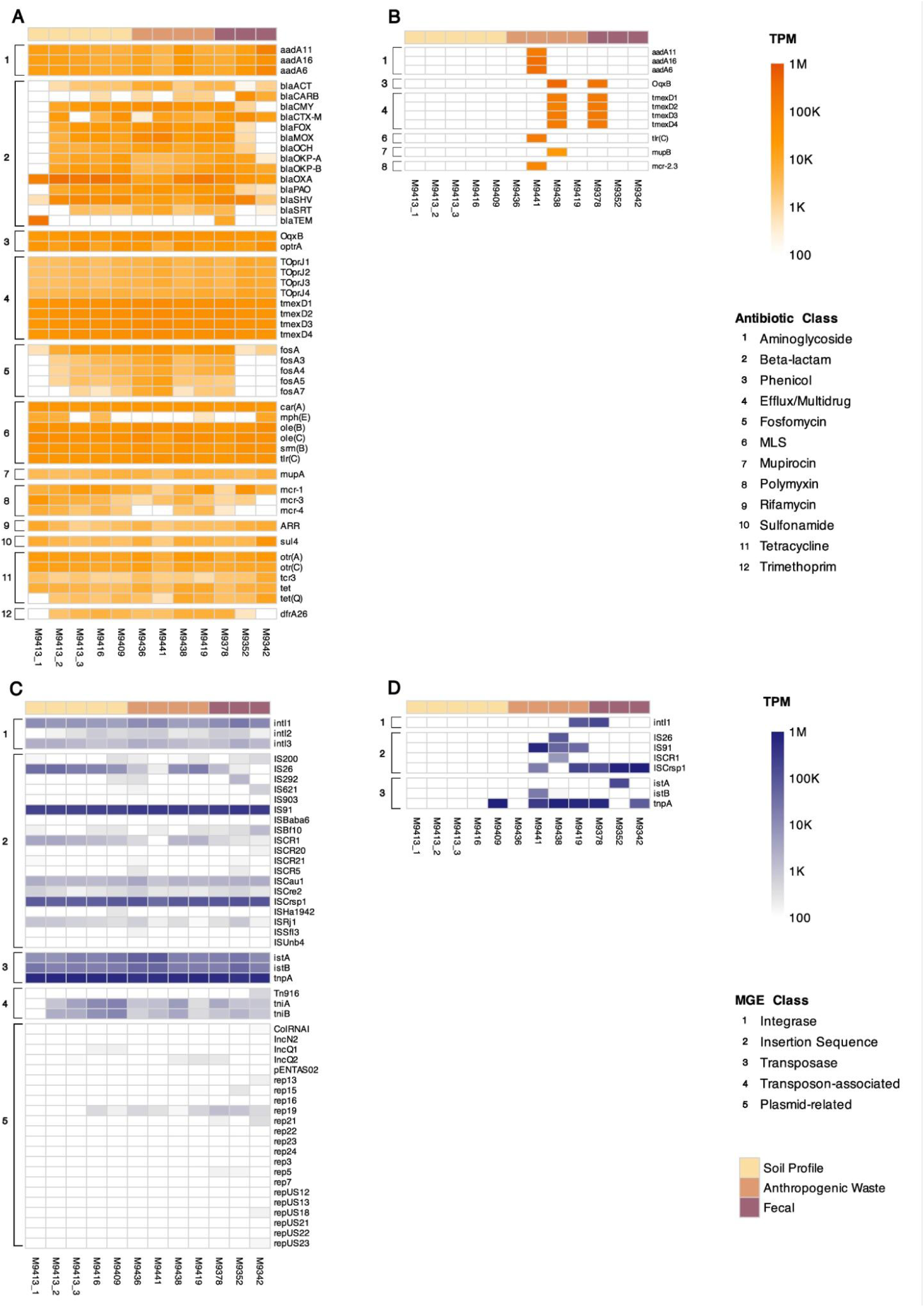
Heatmaps showing antibiotic resistance genes (ARGs) and mobile genetic elements (MGEs) detected across 12 archaeological samples using BLASTN against the ResFinder database (Ferrer Florensa et al. 2022) and a custom MGE database (Pärnänen et al. 2018), respectively. Panels A and C show results from the entire metagenomic dataset, while reads mapped to pyDamage-authenticated ancient DNA contigs are visualized in panels B and D. (A) The 50 most abundant ARG families in the entire metagenomic dataset, ordered by resistance class. (B) All ARG families detected in the damaged dataset, ordered by resistance class. (C) The 50 most abundant MGE families in the entire metagenomic dataset, ordered by MGE class. (D) All MGE families detected in the damaged dataset, ordered by MGE class. The three colours at the top of each heatmap indicate the archaeological sample material type: Soil Profile, Anthropogenic Waste, and Fecal. Samples are displayed on the x-axis and detected gene families on the y-axis. BLASTN hits were retained at a percentage identity ≥ 80%, alignment length ≥ 40 bp, and bitscore ≥ 50. Read counts were normalized using Transcripts Per Million (TPM), and colour scales represent TPM values of 100–1,000,000. Prior to visualization, sequence variants of the same gene were grouped into gene families by removing the variant suffixes, and TPM values were calculated for the variants within each gene family. ARGs are grouped by target antibiotic class, with MLS denoting macrolide-lincosamide-streptogramin. MGEs are grouped into five classes: Integrase, Insertion Sequence, Transposase, Transposon-associated, and Plasmid-related. White cells indicate absence of detection.

A total of 4,071,062 mobile genetic element (MGE) hits were detected after mapping all metagenomic reads against a custom MGE database (Pärnänen et al., 2018) using nucleotide BLAST. After quality filtering, truncating gene variant suffixes, and merging counts of identical gene families, 82 distinct MGE families remained, of which the 50 most abundant are visualized in a heatmap (Fig. 4C). MGEs were classified into five categories: plasmid-related elements (56.10%), insertion sequences (32.93%), integrases (3.66%), transposases (3.66%), and transposon-associated elements (3.66%). The 32 MGE families not shown in the top 50 heatmap comprised plasmid-related elements (n=24) and insertion sequences (n=8), and like many of the displayed plasmid-related hits, these fell below the colour scale minimum of 100 TPM (Supplementary Fig. 2). Across all samples, the ten most abundant MGE families in descending order were *tnpA*, IS*91*, IS*Crsp1*, *istB*, *istA*, IS*26*, *intI1*, *tniA*, *tniB*, and IS*Cau1*. An exception was observed for soil sample M9413_1, representing very low human impact, where only three hits to *tniA* and one hit to *tniB* were detected. Overall, transposases were the most abundant MGE class, while plasmid-related elements were distinctively less abundant. No clear patterns in MGE distribution were observed with respect to the primary material type of the archaeological samples.

Also, the damaged reads were mapped against the ResFinder database (Ferrer Florensa et al., 2022) and MGE database (Pärnänen et al., 2018) using the same BLASTN-based approach applied to the complete metagenomic dataset, and a total of 10,693 MGE hits and 796 ARG hits were recovered. Following quality filtering, 419 ARG hits remained in the damaged dataset, corresponding to 11 unique ARGs detected across three of the twelve samples (M9438, M9378, and M9441; Fig 4B). The detected genes, listed in the order of summed relative abundance, were *oqxB, tmexD4, tmexD3, tmexD2, tmexD1, mupB, mcr*-*2.3, aadA6, aadA16, aadA11,* and *tlr(C)*.

Prior to quality filtering, sample M9419 also yielded hits to *tmexD1–tmexD4* genes with a percent identity of 76.59%, which is below the applied cutoff threshold of 80% and was therefore excluded from downstream analyses and not visualized in the heatmap (Fig. 4B). Notably, all 11 ARGs detected in damaged metagenomes were exclusively from samples originating from a garbage pit (M9438) and a sample containing ash taken from inside the house (M9441), or from a sample of fecal material from the barn floor layer (M9378) (Fig 4B).

ARGs were additionally screened directly from the damaged contigs using the resistance gene identifier (RGI) against the CARD database. However, this yielded only a single hit at a strict cutoff: fosfomycin thiol transferase (*FosI)*, detected in a damaged contig from sample M9441 originating from a sample containing ash taken from inside the house (percent identity 73.08%, bitscore 200).

Quality filtering of the MGE BLAST results yielded 6,742 hits. After removal of gene variant suffixes and merging of identical MGE families, eight unique MGE families were retained: ISCrsp1, *tnpA*, *istA*, *intI1*, IS91, ISCR1, *istB*, and IS26 (Fig. 4D). High-confidence MGE matches (percent identity ≥ 80%, alignment length ≥ 40 bp, bitscore ≥ 50) were detected in seven of the twelve samples. Notably, all soil samples lacked MGE hits, with the exception of sample M9409, collected from the slaughterhouse yard, which contained a hit to *tnpA*. The remaining MGE hits were detected exclusively in samples containing waste or fecal material, following a pattern similar to that observed for ARGs. In addition, the integrase *intI1* was detected in samples collected from the waste pit inside the barn (M9419) and from manure sampled in the barn (M9378).

### Microbial communities reveal associations between resistome and mobilome

Associations between microbial communities, the resistome, and mobile genetic elements were tested using the entire metagenomic dataset, as it is not possible to determine whether the DNA originates from preserved spores, living cells including potential modern contaminants, or extracellular DNA that may be ancient. ARG composition showed significant correlation with microbial community structure at the species level (Mantel test r = 0.744, Benjamini-Hochberg-adjusted p-value, p(BH) < 0.01; PROTEST R² = 0.827, p(BH) < 0.01) (Fig. 5A), indicating that samples with similar ARG profiles also harboured similar microbial communities. MGE composition showed a modest correlation with the microbial community in the Mantel test (r = 0.418, p(BH) = 0.018), while PROTEST indicated a slightly higher agreement (R² = 0.585, p(BH) = 0.031) (Fig. 5B). With resistome and mobilome, the ARG profiles were explained by MGE profiles with both the Mantel test (r = 0.360, p(BH) = 0.030) and PROTEST (R² = 0.689, p(BH) < 0.01) (Fig. 4C). In this analysis, more variance was explained by PROTEST possibly due to some samples having a strong agreement (Fig. 5C) and Mantel’s test analyses the overall agreement of distance matrices.

**Fig 5.**
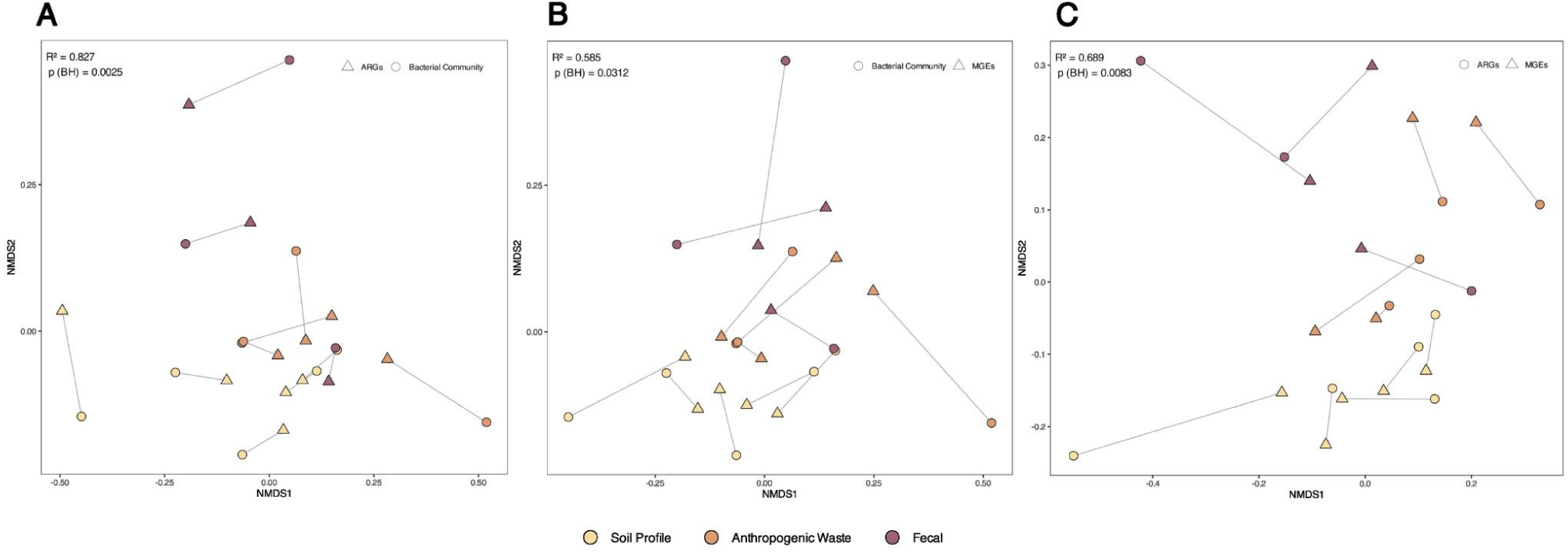
Procrustes analysis analyzing the compositional structure of ARG profiles, MGE profiles, and microbial communities across 12 archaeological metagenomic samples. Each plot shows the ordination of two datasets after symmetric Procrustes superimposition of their NMDS configurations based on Hellinger-transformed Bray-Curtis dissimilarities. Prior to ordination, ARG and MGE sequence variants were collapsed into canonical gene families by stripping trailing variant suffixes and summing Transcripts Per Million (TPM) values across variants within each family. Arrows connect each sample’s position between the two ordinations, with shorter arrows indicating greater agreement between community structures. The PROTEST correlation coefficient (R²) indicates the degree of similarity between the two ordinations, and the Benjamini-Hochberg adjusted permutation p-value (p(BH); 999 permutations) indicates statistical significance. A) ARGs and microbial community composition at genus level. B) MGEs and microbial community composition at genus level. C) ARGs and MGEs.

### Waste- and fecal-associated samples yielded diverse Gram-Positive isolates including potentially novel taxa

A total of 14 Gram-positive bacterial isolates were recovered, representing 11 distinct species (Table 2). Only one isolate, putative *Arthrobacter sp*. (M9409_L16), was recovered from a soil sample exposed to high human impact (House yard). The remaining isolates were recovered from samples containing wastes (n = 7) or fecal materials (n = 6). High-quality genomes were obtained across all the isolates, with genome completeness values varying between 99.97–100% and contamination between 0.03–5.37%. The highest estimated contamination level of 5.37% was observed in *Paeniglutamicibacter sulfureus* (M9419_N6). Two isolates, putative *Arthrobacter sp*. (M9409_L16) and putative *Psychrobascillus sp*. (M9378_M5), showed average nucleotide identity (ANI) values of 92.99% and 92.84%, respectively, indicating that they could be novel species. Putative *Nocardioides sp*. (M9419_N2) exhibited an ANI value of 86.32%, which could potentially represent a novel genus. All remaining isolates had ANI values above 97%, indicating reliable taxonomical classification.

**Table 2.**
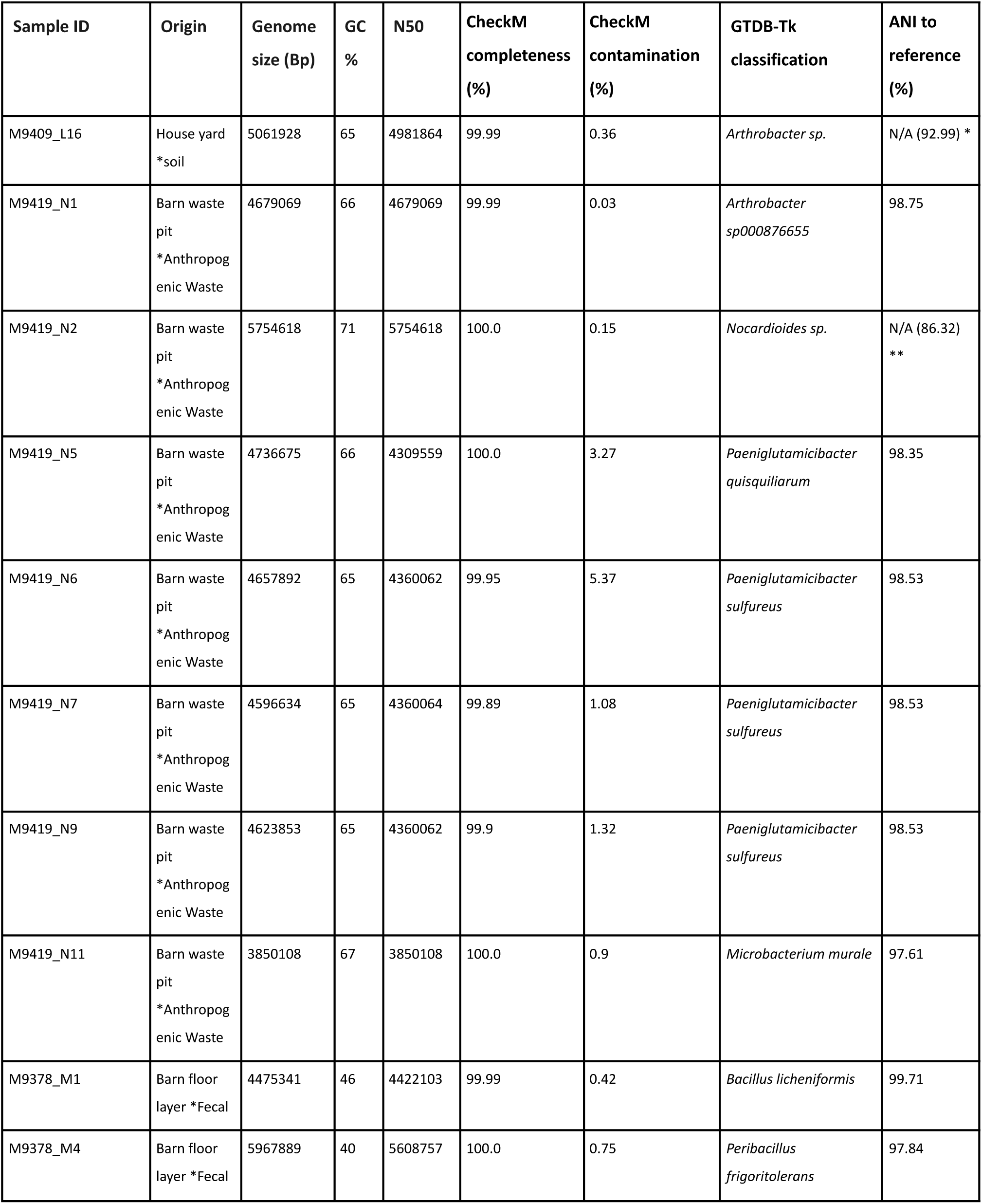
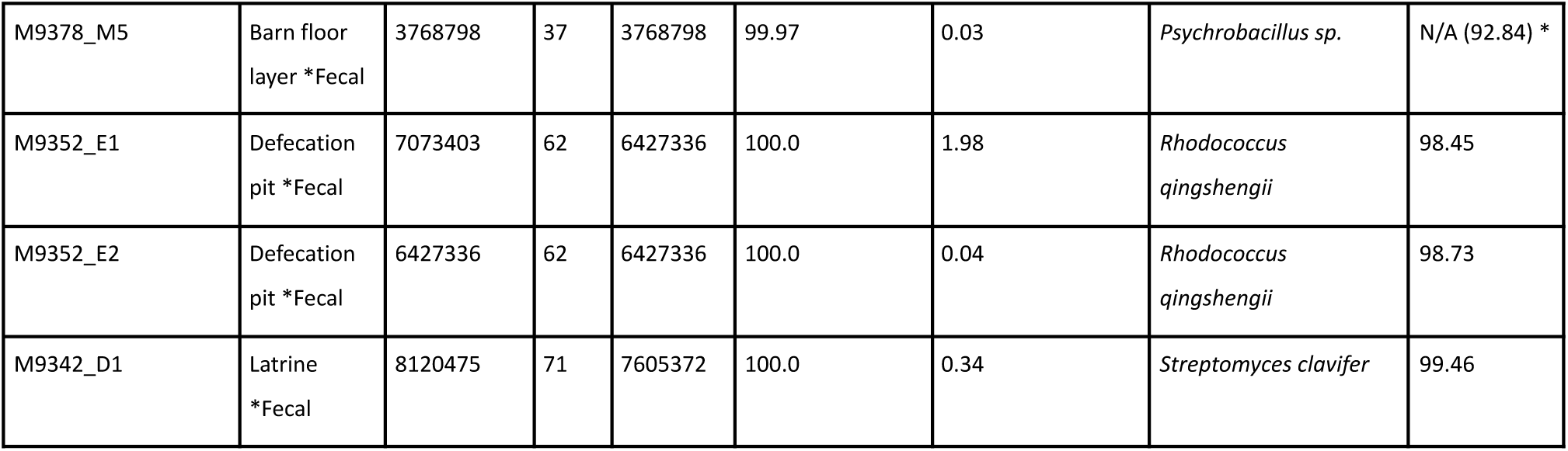
Genome assembly statistics and taxonomic classification of gram-positive bacterial isolates derived from archaeological samples. * Isolate falls outside the average nucleotide identity (ANI) radius of the closest reference species, indicating a putative novel species. ** No reference genome exceeded the ANI calculation threshold; no formal species assignment could be made. For both * and **, the closest placement ANI is reported in parentheses.

All isolate genomes were detected in each metagenomic sample following read mapping (Supplementary Fig. 3). A putative *Psychrobacillus sp*. was the most abundant isolate in samples containing anthropogenic waste (M9419) and fecal material (M9352 and M9342), even though it was isolated from sample M9378 (Barn floor). The abundances of the three *P. sulfureus* isolates and the two *Rhodococcus qingshengii* isolates were similar across all metagenomic samples. None of the isolates were among the most abundant in that sample it originated from. Overall, all isolates were clearly less abundant in soil samples reflecting the microbiomes before urbanization (M9413_1–3) than in samples reflecting the 1800 century slaughterhouse environment (Supplementary Fig. 3).

### Isolates harbor multiple antibiotic resistance genes, mobile genetic elements, and biosynthetic gene clusters encoding antimicrobial activity

A total of seven antibiotic resistance genes (ARGs) and five mobile genetic elements (MGEs) were detected across the genomes of the isolates after quality filtering with a minimum threshold of 70% identity and ≥40% coverage, retaining only the best hit per genomic location (Fig. 6). The ARGs were categorized into six antibiotic classes: Beta-lactam (*blaZ*), Chloramphenicol (*cat* and *cmlV)*, Macrolide (*ole(C)*), Macrolide-Lincosamide-Streptogramin (*erm(D)*), Rifamycin (*ARR-7*), and Tetracycline (*tet(43)*). The MGEs were classified into four types: insertion sequence (IS*Cau1*), integrase (*int2*), transposase (*tnpA*), and transposon-associated proteins (*tniA* and *tniB*).

**Fig 6.**
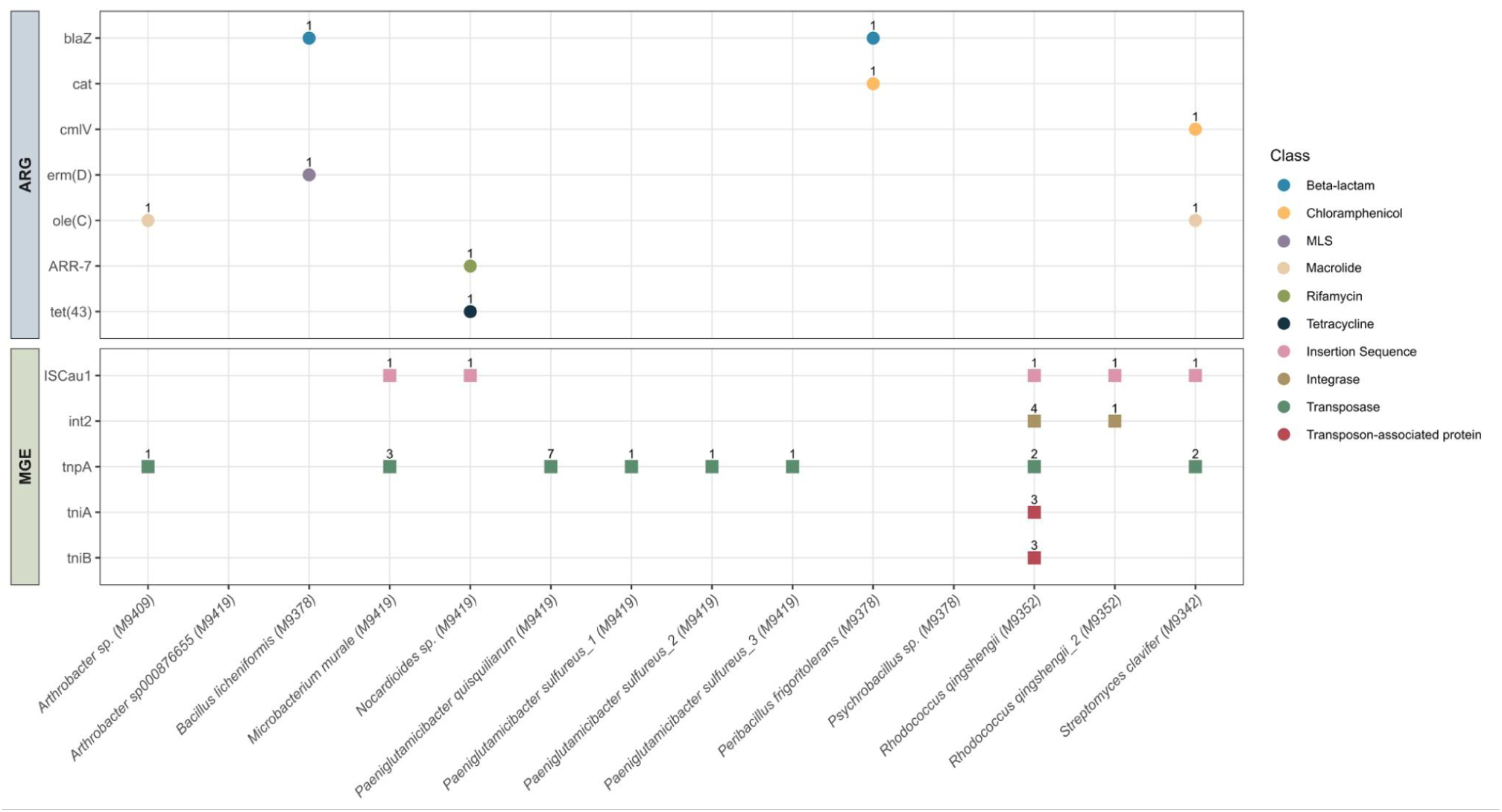
Dot plot of antibiotic resistance genes (ARGs) and mobile genetic elements (MGEs) detected in 14 Gram-positive isolates from 18th century archaeological samples. ARGs were identified by nucleotide BLAST against the ResFinder database (≥70% identity, ≥60% coverage) and MGEs against the database of Pärnänen et al. (2018) (≥70% identity, ≥40% coverage). If multiple hits overlapped on the same contig, only the best hit by percent identity was retained. Numbers above symbols indicate the count of the detected gene. Color of the symbol indicates antibiotic class or mobile genetic element class. The isolate name based on the GTDB-Tk classification is followed by the archaeological sample code in parentheses.

Among the actinobacterial isolates, the putative *Arthrobacter sp.* carried both *ole(C)* and tnpA, while *Arthrobacter sp000876655* had no detectable ARGs or MGEs. *Nocardioides sp.* harboured two ARGs, *ARR-7* and *tet(43)*, alongside IS*Cau1*. *Microbacterium murale* contained no ARGs but carried one IS*Cau1* and three *tnpA* genes. *Paeniglutamicibacter quisquiliarum* lacked ARGs but had seven *tnpA* genes distributed across different genomic regions, while none of the three *P. sulfureus* isolates contained ARGs, and each of the isolates was also carrying a single *tnpA* gene. The two *R. qingshengii* isolates differed in their MGE profiles: one carried IS*Cau1*, four *int2*, two *tnpA* genes, and three copies each of *tniA* and *tniB,* while the other had only a single copy of IS*Cau1* and *int2*; neither isolate harboured ARGs. *Streptomyces clavifer* harbored the resistance genes *cmlV* and *ole(C)*, together with the MGEs IS*Cau1* and two copies of *tnpA*. The genomic locations of *cmlV* and IS*Cau1* were relatively close to each other, with *cmlV* located between positions 3,424,892 and 3,425,991, and IS*Cau1* between positions 3,875,823 and 3,876,457 (Supplementary Table 3). Among the *Bacillales* isolates, the *blaZ* gene was detected in both *Bacillus licheniformis* and *Peribacillus frigoritolerans*. *B. licheniformis* additionally harbored *erm(D)* but did not contain any detectable MGEs. In *P. frigoritolerans*, *blaZ* co-occurred with the *cat* gene, and the two resistance genes were located relatively close to each other in the genome, with *blaZ* located between positions 5,305,415 bp and 5,305,983 bp and *cat* between positions 5,364,656 bp and 5,365,267 bp. No MGEs were detected in this isolate. Putative *Psychrobacillus* sp. did not contain any ARGs or MGEs.

A total of 2,986 MIBiG (Minimum Information about a Biosynthetic Gene Cluster) entries were parsed, of which 769 were associated with antibiotic production. After filtering for hits with more than 15% similarity to known antibiotic-producing clusters, 16 hits were retained across the Gram-positive isolates (Fig. 7). The most confident predictions were those with 100% similarity to known biosynthetic gene clusters (BGCs). *B. licheniformis* harboured three such hits: lichenysin, lichenicidin VK21 A1/A2, and bacillibactin E/F, alongside a 66% similarity hit to pulcherriminic acid. A BGC with 100% similarity to ε-poly-L-lysine was identified in the putative *Nocardioides sp.* and in both *R. qingshengii* isolates. The two *R. qingshengii* isolates additionally carried a BGC with 93% similarity to corynecin I/II/III and a 57% similarity hit to erythrochelin. A BGC with 31% similarity to stenothricin was detected in the putative *Arthrobacter sp.* and *Arthrobacter sp000876655*, with related hits at 22% similarity in *P. quisquiliarum* and 27% similarity in *P. sulfureus. S. clavifer* contained four BGCs with low similarity to known antibiotic clusters: hits to steffimycin D and kinamycin at 16%, a BGC with 20% similarity to clavulanic acid, and a BGC with 24% similarity to argimycin PI/PII/PIV/PV/PVI/PIX and nigrifactin.

**Fig 7.**
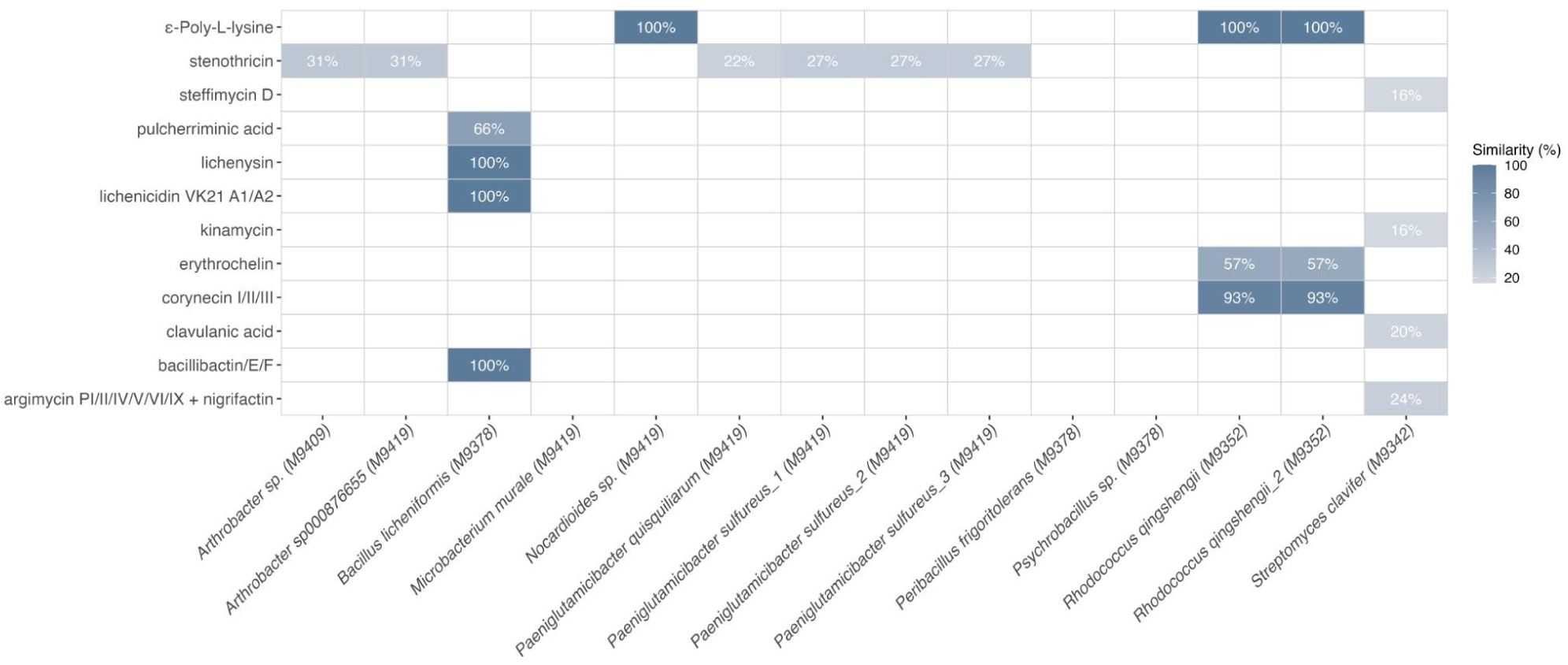
Heatmap of predicted biosynthetic gene cluster (BGC) similarity to known antibiotic-producing reference clusters from the Minimum Information about a Biosynthetic Gene Cluster database (MIBiG). BGC prediction was performed with antiSMASH, and only BGCs with ≥15% similarity to a reference cluster annotated with antibacterial activity in MIBiG were retained. The x-axis shows the isolate names (GTDB-Tk classification) followed by the archaeological sample code in parentheses and the y-axis shows MIBiG reference cluster names. Cell color and the percentage printed within each cell indicate the KnownClusterBlast similarity score between the predicted BGC and its closest MIBiG reference.

## Discussion

To illuminate the foundations of the AMR problem, we combined both metagenomic and culture-based approaches to study microbiomes in the 18th-century urban environment with slaughterhouse activity. Rather than focusing on individual organisms, we applied a holistic approach to obtain an overview of pre-industrial, pre-antibiotic-era microbial community structures, resistomes, and mobilomes. The top 15 most abundant bacterial orders and the top 50 most abundant genera in the metagenomic dataset predominantly represented taxa commonly found in environmental samples. The antibiotic resistance gene (ARG) profile observed in this study reflects a typical soil-associated resistome (D’Costa et al., 2011b; Gibson et al., 2015; Martínez, 2008), with genes conferring resistance to antibiotics produced by environmental microorganisms. These included resistance to beta-lactams, aminoglycosides, Macrolide–Lincosamide–Streptogramins (MLS), tetracyclines, rifamycins, glycopeptides, and fusidic acid, as well as to synthetic antibiotics such as quinolones, sulfonamides, trimethoprim, phenicols, fosfomycin, and mupirocin.

We were able to establish that antibiotic resistance may have begun to disseminate already in pre-industrial urban environments as we detected ARGs, MGEs and recovered dormant antibiotic-producing bacteria in samples from anthropogenic wastes. The waste pits and piles were favorable habitats for antibiotic-producing organisms since they were rich in organic matter and in direct contact with soil. The wastes also received constant inputs of human- and livestock-derived fecal materials containing gut bacteria known for carrying MGEs (Wibowo et al., 2021b). Antibiotic producers most likely created selection pressures that fostered the integration of genes of different bacteria into the genomes of one another, including ARGs and MGEs (Gillings and Stokes, 2012b; Peterson and Kaur, 2018b; Wellington et al., 2013). This interpretation is supported by the Procrustes and Mantel test results, as well as the genome analysis of dormant Gram-positive bacteria carrying MGEs commonly found in fecal-associated bacteria, in addition to their ARGs and antibiotic-production-related biosynthetic gene clusters.

Intriguingly, our analysis revealed the presence of homologous genes encoding the Resistance-Nodulation-Cell Division (RND) -type efflux pump system (TmexCD-TOprJ), across the entire metagenomic dataset, and *tmexD1-4* genes were detectable within the damaged metagenomes. The *tmexC1-4*, *tmexD1-4* and *TOprJ1-4* genes are components of a plasmid-borne RND-family efflux pump system, unique in Gram-negative bacteria and found in strains of Enterobacteriaceae and Pseudomonas (Dong et al., 2022a; Gao et al., 2022). RND efflux pumps are typically encoded chromosomally, but this novel plasmid-borne efflux pump, tmexCD1-TOprJ1, was first reported in 2020 (Lv et al., 2020). These pumps confer resistance to multiple classes of antibiotics such as tetracyclines, including the last-resort antibiotic tigecycline. There have been indications that these genes potentially originate from *Pseudomonas* spp. (Dong et al., 2022b; Lv et al., 2020; Wang et al., 2021). In our results, *Pseudomonas* was the most abundant genera in the entire metagenomic dataset and second most abundant in the damaged metagenomic dataset. *Pseudomonas* spp. are ubiquitous bacteria inhabiting diverse ecological niches and include species known to harbor intrinsic antibiotic resistance mechanisms that can be transferred via horizontal gene transfer to clinically relevant strains (Luczkiewicz et al., 2015; Mohanty et al., 2026).

We also analyzed the contigs assembled from aDNA reads, and detected the fosfomycin thiol transferase gene *fosI*, which confers resistance to the antibiotic fosfomycin, introduced in the 1960s and is widely used in the treatment of multidrug-resistant infections. It is known to be produced by some *Streptomycetes* and *Pseudomonas* species (Hendlin et al., 1969; Simon et al., 2021) and was first described as an integron-associated gene cassette (Pelegrino et al., 2015). The damaged contig containing the *fosI* gene was recovered from a sample containing ash collected from a kitchen from a house in the courtyard. Fosfomycin resistance gene families have previously been reported from clinical pathogens as well as from diverse environmental settings, including wastewater, sediments, and other anthropogenically impacted habitats (Ito et al., 2017; Kieffer et al., 2025). Kieffer et al., 2025 suggested that environmental contamination selects fosfomycin resistance genes and this is supported by our detection as ash is extremely alkaline and may contain heavy metals and thus creates significant selection pressure for microbiomes.

All cultivated isolates originated from samples containing waste or fecal material. With the exception of *Psychrobacillus* sp., all isolates possessed either ARGs, MGEs, BGCs encoding antimicrobial compounds, or a combination of these. *Bacillus licheniformis* was isolated from a barn sample, from a manure-rich layer composed of straw and decomposed organic material that likely derived from livestock. This isolate carried the beta-lactam (*blaZ*) and Macrolide–Lincosamide–Streptogramin (*erm*(D)) resistance genes. While *erm*(D) has previously been reported in *B. licheniformis* and may constitute part of the species’ ancient resistome (Agersø et al., 2019), *bla*Z has not to our knowledge been reported previously in this species. In addition, the strain harbored several antibiotic-production-associated BGCs, including lichenysin, lichenicidin VK21 A1/A2, and pulcherriminic acid, all previously described in *B. licheniformis* (Caetano et al., 2011; Wang et al., 2018; Yakimov et al., 1999), as well asbacillibactin E/F cluster, which has previously been reported from *B. haynesii* rather than *B. licheniformis* (Eltokhy et al., 2024).

In metagenomic analyses of the barn sample, the damaged dataset included taxa of both environmental and gut-associated origin, some of which contain pathogenic species, such as *Campylobacter*, *Streptococcus* and *Helicobacter*, in addition to copiotrophic bacteria such as *Pseudomonas*, *Bacillus*, *Streptomyces*, and *Paenibacillus*, which thrive in nutrient-rich environments containing organic matter and produce antibiotics (Lee et al., 2023; Olishevska et al., 2019). We also detected *intl1* in the aDNA reads of this sample, and in one additional fecal material containing sample, as wel as in all samples across the entire metagenomic dataset. *Intl1* can be considered as an indicator of anthropogenic pollution, as it is linked to ARGs, disinfectants, and heavy metals; all of which can be transferred through horizontal gene transfer (Gillings et al., 2015; Gillings and Stokes, 2012a; Wellington et al., 2013). Together, the detection of potential pathogenic bacteria, antibiotic producing copiotrophs and *intI1* in metagenomic data further supports our hypothesis that preindustrial waste- and manure-rich environments acted as ecological hotspots for genetic exchange between environmental, gut-associated, and potentially pathogenic bacteria.

Due to the high microbial diversity in the metagenomic data, aDNA analysis was unsuccessful using methods commonly applied to aDNA reads (Jónsson et al., 2013; Pochon et al., 2023). To enable read-based analyses of microbiomes, ARGs, and MGEs, we generated metagenomic contigs and authenticated those containing damaged aDNA using pyDamage (Borry et al., 2021). Mapping all reads against the authenticated contigs allowed us to separate the damaged read fraction and analyze its taxonomy, resistome, and mobilome. However, the analysis of resistomes and mobilomes remained challenging. The pyDamage-based aDNA authentication approach requires metagenomic assemblies, which are known to represent a bottleneck for ARG and MGE detection, as these elements are often poorly assembled due to their diverse genetic contexts and environments (Abramova et al., 2024; Kerkvliet et al., 2024). Assembly of soil metagenomes has also been shown to be challenging due to the complexity and biodiversity of the soil community, as well as gaps in the reference databases (Curtis et al., 2002; Fierer et al., 2007). Nevertheless, due to the limited availability of aDNA authentication tools suitable for samples with such high bacterial diversity, pyDamage represented the most suitable approach.

In DNA extraction, we used a column-free method and magnetic beads, which made it possible to capture both short and long DNA fragments, enabling the recovery of short and fragmented DNA (aDNA) as well as DNA in viable and dormant cells. Due to the relatively young age of the archaeological site and the potential contribution of dormant bacterial spores to our research question, it was meaningful to study both DNA fractions, i.e. DNA derived from both dormant cells and highly fragmented aDNA released by dead cells. Therefore we used ARG and MGE post-filtering thresholds (≥80% identity, ≥40 bp alignment length, ≥50 bitscore) that resulted in a considerable number of relatively short hits. Thus, some of the detected ARG hits should be interpreted with caution. Also, due to the relatively young age of the archaeological site, radiocarbon dating was not feasible. However, archaeological findings (Uotila et al., 2025) and stratigraphic layers associated with urban fires confirm that the samples originate from the 18th century.

This study demonstrated that environmental archaeological samples containing different types of anthropogenic wastes provide an beyond state-of-the-art approach for investigating the microbiome, resistome, and mobilome in preindustrial human impacted environments. Holistic approaches like this could for instance, deliver new understanding on bacterial adaptations to human influenced environmental changes, which could, in turn, yield discoveries of alternatives for antibiotics. Although it now seems our race against antibiotic resistance might have become impossible to win, we might find alternative approaches through understanding the early evolution and ecological history of AMR.

## Data availability

Metagenomic sequencing data will be made available upon publication. Filtered sequencing data and genomes assemblies of isolate genomes are available in the European Nucleotide Archive (ENA) under accession number PRJEB114365. Bioinformatic pipelines for metagenomic and whole genomic analysis and data visualization R scripts will be made available upon publication.

## Author contributions

Minna Maria Maunula: Conceptualization, Data curation, Formal analysis, Investigation, Methodology, Project administration, Visualization, Writing – original draft, and Writing – review and editing. Taru-Marja Mäkinen: Investigation, Methodology, Software, Validation, Writing – Review & Editing. Kari Uotila: Investigation, Resources and Writing – review and editing. Kirill Bodganov: Investigation and Writing – review and editing. Marko Virta: Supervision, Funding acquisition, Resources, and Writing – review and editing. Jenni Hultman: Conceptualization, Funding acquisition, Methodology, Project administration, Resources, Supervision, and Writing – review and editing. Johanna Muurinen: Conceptualization, Funding acquisition, Methodology, Project administration, Resources, Supervision, and Writing – review and editing.

## Use of Artificial Intelligence

To improve the grammar, readability, and overall clarity of the manuscript, Microsoft 365 Copilot (Microsoft, Redmond, WA, USA) was used for language editing and refinement. Claude Sonnet 4.6 (Anthropic, San Francisco, CA, USA) was used to generate and improve the computational code. All generated material was reviewed, revised, and verified by the authors. Scientific interpretation, analysis of findings, and conclusions were performed exclusively by the authors.

## Acknowledgements

We acknowledge the DNA Sequencing and Genomics Laboratory at the Institute of Biotechnology, University of Helsinki, supported by HiLIFE and Biocenter Finland, for sequencing services.

Computational resources were provided by CSC – IT Center for Science, Finland, through the Puhti computing environment.

## References

Abramova, A., Karkman, A., Bengtsson-Palme, J., 2024. Metagenomic assemblies tend to break around antibiotic resistance genes. BMC Genomics 25, 959. 10.1186/s12864-024-10876-0

Agersø, Y., Bjerre, K., Brockmann, E., Johansen, E., Nielsen, B., Siezen, R., Stuer-Lauridsen, B., Wels, M., Zeidan, A.A., 2019. Putative antibiotic resistance genes present in extant Bacillus licheniformis and Bacillus paralicheniformis strains are probably intrinsic and part of the ancient resistome. PLOS ONE 14, e0210363. 10.1371/journal.pone.0210363

Altschul, S.F., Gish, W., Miller, W., Myers, E.W., Lipman, D.J., 1990. Basic local alignment search tool. J. Mol. Biol. 215, 403–410. 10.1016/S0022-2836(05)80360-2

Berkner, S., Konradi, S., Schönfeld, J., 2014. Antibiotic resistance and the environment—there and back again. EMBO Rep. 15, 740–744. 10.15252/embr.201438978

Berthold, T., Centler, F., Hübschmann, T., Remer, R., Thullner, M., Harms, H., Wick, L.Y., 2016. Mycelia as a focal point for horizontal gene transfer among soil bacteria. Sci. Rep. 6, 36390. 10.1038/srep36390

Blin, K., Shaw, S., Vader, L., Szenei, J., Reitz, Z.L., Augustijn, H.E., Cediel-Becerra, J.D.D., de Crécy-Lagard, V., Koetsier, R.A., Williams, S.E., Cruz-Morales, P., Wongwas, S., Segurado Luchsinger, A.E., Biermann, F., Korenskaia, A., Zdouc, M.M., Meijer, D., Terlouw, B.R., van der Hooft, J.J.J., Ziemert, N., Helfrich, E.J.N., Masschelein, J., Corre, C., Chevrette, M.G., van Wezel, G.P., Medema, M.H., Weber, T., 2025. antiSMASH 8.0: extended gene cluster detection capabilities and analyses of chemistry, enzymology, and regulation. Nucleic Acids Res. 53, W32–W38. 10.1093/nar/gkaf334

Borry, M., Hübner, A., Rohrlach, A.B., Warinner, C., 2021. PyDamage: automated ancient damage identification and estimation for contigs in ancient DNA de novo assembly. PeerJ 9, e11845. 10.7717/peerj.11845

Caetano, T., Krawczyk, J.M., Mösker, E., Süssmuth, R.D., Mendo, S., 2011. Heterologous Expression, Biosynthesis, and Mutagenesis of Type II Lantibiotics from *Bacillus licheniformis* in *Escherichia coli*. Chem. Biol. 18, 90–100. 10.1016/j.chembiol.2010.11.010

Cantón, R., 2009. Antibiotic resistance genes from the environment: a perspective through newly identified antibiotic resistance mechanisms in the clinical setting. Clin. Microbiol. Infect. 15, 20–25. 10.1111/j.1469-0691.2008.02679.x

Carini, P., Marsden, P.J., Leff, J.W., Morgan, E.E., Strickland, M.S., Fierer, N., 2016. Relic DNA is abundant in soil and obscures estimates of soil microbial diversity. Nat. Microbiol. 2, 16242. 10.1038/nmicrobiol.2016.242

Chaumeil, P.-A., Mussig, A.J., Hugenholtz, P., Parks, D.H., 2022. GTDB-Tk v2: memory friendly classification with the genome taxonomy database. Bioinformatics 38, 5315–5316. 10.1093/bioinformatics/btac672

Chen, S., Zhou, Y., Chen, Y., Gu, J., 2018. fastp: an ultra-fast all-in-one FASTQ preprocessor. Bioinformatics 34, i884–i890. 10.1093/bioinformatics/bty560

Cheng, H., Concepcion, G.T., Feng, X., Zhang, H., Li, H., 2021. Haplotype-resolved de novo assembly using phased assembly graphs with hifiasm. Nat. Methods 18, 170–175. 10.1038/s41592-020-01056-5

Chklovski, A., Parks, D.H., Woodcroft, B.J., Tyson, G.W., 2023. CheckM2: a rapid, scalable and accurate tool for assessing microbial genome quality using machine learning. Nat. Methods 20, 1203–1212. 10.1038/s41592-023-01940-w

Courvalin, P., 2008. Predictable and unpredictable evolution of antibiotic resistance. J. Intern. Med. 264, 4–16. 10.1111/j.1365-2796.2008.01940.x

Curtis, T.P., Sloan, W.T., Scannell, J.W., 2002. Estimating prokaryotic diversity and its limits. Proc. Natl. Acad. Sci. U. S. A. 99, 10494–10499. 10.1073/pnas.142680199

Danecek, P., Bonfield, J.K., Liddle, J., Marshall, J., Ohan, V., Pollard, M.O., Whitwham, A., Keane, T., McCarthy, S.A., Davies, R.M., Li, H., 2021. Twelve years of SAMtools and BCFtools. GigaScience 10, giab008. 10.1093/gigascience/giab008

D’Costa, V.M., King, C.E., Kalan, L., Morar, M., Sung, W.W.L., Schwarz, C., Froese, D., Zazula, G., Calmels, F., Debruyne, R., Golding, G.B., Poinar, H.N., Wright, G.D., 2011a. Antibiotic resistance is ancient. Nature 477, 457–461. 10.1038/nature10388

D’Costa, V.M., King, C.E., Kalan, L., Morar, M., Sung, W.W.L., Schwarz, C., Froese, D., Zazula, G., Calmels, F., Debruyne, R., Golding, G.B., Poinar, H.N., Wright, G.D., 2011b. Antibiotic resistance is ancient. Nature 477, 457–461. 10.1038/nature10388

De Simeis, D., Serra, S., 2021. Actinomycetes: A Never-Ending Source of Bioactive Compounds—An Overview on Antibiotics Production. Antibiotics 10, 483. 10.3390/antibiotics10050483

DeAngelis, K.M., Brodie, E.L., DeSantis, T.Z., Andersen, G.L., Lindow, S.E., Firestone, M.K., 2009. Selective progressive response of soil microbial community to wild oat roots. ISME J. 3, 168–178. 10.1038/ismej.2008.103

Dong, N., Zeng, Y., Wang, Yao, Liu, C., Lu, J., Cai, C., Liu, X., Chen, Y., Wu, Yuchen, Fang, Y., Fu, Y., Hu, Y., Zhou, H., Cai, J., Hu, F., Wang, S., Wang, Yang, Wu, Yongning, Chen, G., Shen, Z., Chen, S., Zhang, R., 2022a. Distribution and spread of the mobilised RND efflux pump gene cluster *tmexCD-toprJ* in clinical Gram-negative bacteria: a molecular epidemiological study. Lancet Microbe 3, e846–e856. 10.1016/S2666-5247(22)00221-X

Dong, N., Zeng, Y., Wang, Yao, Liu, C., Lu, J., Cai, C., Liu, X., Chen, Y., Wu, Yuchen, Fang, Y., Fu, Y., Hu, Y., Zhou, H., Cai, J., Hu, F., Wang, S., Wang, Yang, Wu, Yongning, Chen, G., Shen, Z., Chen, S., Zhang, R., 2022b. Distribution and spread of the mobilised RND efflux pump gene cluster *tmexCD-toprJ* in clinical Gram-negative bacteria: a molecular epidemiological study. Lancet Microbe 3, e846–e856. 10.1016/S2666-5247(22)00221-X

Edwin, N.R., Fitzpatrick, A.H., Brennan, F., Abram, F., O’Sullivan, O., 2024. An in-depth evaluation of metagenomic classifiers for soil microbiomes. Environ. Microbiome 19, 19. 10.1186/s40793-024-00561-w

Eltokhy, M.A., Saad, B.T., Eltayeb, W.N., Alshahrani, M.Y., Radwan, S.M.R., Aboshanab, K.M., Ashour, M.S.E., 2024. Metagenomic nanopore sequencing for exploring the nature of antimicrobial metabolites of Bacillus haynesii. AMB Express 14, 52. 10.1186/s13568-024-01701-8

Ewels, P., Magnusson, M., Lundin, S., Käller, M., 2016. MultiQC: summarize analysis results for multiple tools and samples in a single report. Bioinformatics 32, 3047–3048. 10.1093/bioinformatics/btw354

Fierer, N., Breitbart, M., Nulton, J., Salamon, P., Lozupone, C., Jones, R., Robeson, M., Edwards, R.A., Felts, B., Rayhawk, S., Knight, R., Rohwer, F., Jackson, R.B., 2007. Metagenomic and small-subunit rRNA analyses reveal the genetic diversity of bacteria, archaea, fungi, and viruses in soil. Appl. Environ. Microbiol. 73, 7059–7066. 10.1128/AEM.00358-07

Florensa, A.F., Kaas, R.S., Clausen, P.T.L.C., Aytan-Aktug, D., Aarestrup, F.M., 2022. ResFinder – an open online resource for identification of antimicrobial resistance genes in next-generation sequencing data and prediction of phenotypes from genotypes. Microb. Genomics 8, 000748. 10.1099/mgen.0.000748

Fluit, A.C., Schmitz, F.J., 1999. Class 1 Integrons, Gene Cassettes, Mobility, and Epidemiology. Eur. J. Clin. Microbiol. Infect. Dis. 18, 761–770. 10.1007/s100960050398

Gao, X., Wang, C., Lv, L., He, X., Cai, Z., He, W., Li, T., Liu, J.-H., 2022. Emergence of a Novel Plasmid-Mediated Tigecycline Resistance Gene Cluster, tmexCD4-toprJ4, in Klebsiella quasipneumoniae and Enterobacter roggenkampii. Microbiol. Spectr. 10, e01094–22. 10.1128/spectrum.01094-22

Gaze, W.H., Zhang, L., Abdouslam, N.A., Hawkey, P.M., Calvo-Bado, L., Royle, J., Brown, H., Davis, S., Kay, P., Boxall, A.B.A., Wellington, E.M.H., 2011. Impacts of anthropogenic activity on the ecology of class 1 integrons and integron-associated genes in the environment. ISME J. 5, 1253–1261. 10.1038/ismej.2011.15

Ghaly, T.M., Gillings, M.R., n.d. New perspectives on mobile genetic elements: a paradigm shift for managing the antibiotic resistance crisis. Philos. Trans. R. Soc. B Biol. Sci. 377, 20200462. 10.1098/rstb.2020.0462

Gibson, M.K., Forsberg, K.J., Dantas, G., 2015. Improved annotation of antibiotic resistance determinants reveals microbial resistomes cluster by ecology. ISME J. 9, 207–216. 10.1038/ismej.2014.106

Gillings, M.R., Gaze, W.H., Pruden, A., Smalla, K., Tiedje, J.M., Zhu, Y.-G., 2015. Using the class 1 integron-integrase gene as a proxy for anthropogenic pollution. ISME J. 9, 1269–1279. 10.1038/ismej.2014.226

Gillings, M.R., Stokes, H.W., 2012a. Are humans increasing bacterial evolvability? Trends Ecol. Evol. 27, 346–352. 10.1016/j.tree.2012.02.006

Gillings, M.R., Stokes, H.W., 2012b. Are humans increasing bacterial evolvability? Trends Ecol. Evol. 27, 346–352. 10.1016/j.tree.2012.02.006

Griffiths, R.I., Whiteley, A.S., O’Donnell, A.G., Bailey, M.J., 2000. Rapid Method for Coextraction of DNA and RNA from Natural Environments for Analysis of Ribosomal DNA- and rRNA-Based Microbial Community Composition. Appl. Environ. Microbiol. 66, 5488–5491. 10.1128/AEM.66.12.5488-5491.2000

Hendlin, D., Stapley, E.O., Jackson, M., Wallick, H., Miller, A.K., Wolf, F.J., Miller, T.W., Chaiet, L., Kahan, F.M., Foltz, E.L., Woodruff, H.B., Mata, J.M., Hernandez, S., Mochales, S., 1969. Phosphonomycin, a New Antibiotic Produced by Strains of Streptomyces. Science 166, 122–123. 10.1126/science.166.3901.122

Ito, R., Mustapha, M.M., Tomich, A.D., Callaghan, J.D., McElheny, C.L., Mettus, R.T., Shanks, R.M.Q., Sluis-Cremer, N., Doi, Y., 2017. Widespread Fosfomycin Resistance in Gram-Negative Bacteria Attributable to the Chromosomal fosA Gene. mBio 8, e00749–17. 10.1128/mBio.00749-17

Jiang, X., Ellabaan, M.M.H., Charusanti, P., Munck, C., Blin, K., Tong, Y., Weber, T., Sommer, M.O.A., Lee, S.Y., 2017. Dissemination of antibiotic resistance genes from antibiotic producers to pathogens. Nat. Commun. 8, 15784. 10.1038/ncomms15784

Jónsson, H., Ginolhac, A., Schubert, M., Johnson, P.L.F., Orlando, L., 2013. mapDamage2.0: fast approximate Bayesian estimates of ancient DNA damage parameters. Bioinformatics 29, 1682–1684. 10.1093/bioinformatics/btt193

Kang, D.D., Li, F., Kirton, E., Thomas, A., Egan, R., An, H., Wang, Z., 2019. MetaBAT 2: an adaptive binning algorithm for robust and efficient genome reconstruction from metagenome assemblies. PeerJ 7, e7359. 10.7717/peerj.7359

Kerkvliet, J.J., Bossers, A., Kers, J.G., Meneses, R., Willems, R., Schürch, A.C., 2024. Metagenomic assembly is the main bottleneck in the identification of mobile genetic elements. PeerJ 12, e16695. 10.7717/peerj.16695

Kieffer, N., Böhm, M.-E., Berglund, F., Marathe, N.P., Gillings, M.R., Larsson, D.G.J., 2025. Identification of novel FosX family determinants from diverse environmental samples. J. Glob. Antimicrob. Resist. 41, 8–14. 10.1016/j.jgar.2024.12.018

Kjær, K.H., Winther Pedersen, M., De Sanctis, B., De Cahsan, B., Korneliussen, T.S., Michelsen, C.S., Sand, K.K., Jelavić, S., Ruter, A.H., Schmidt, A.M.A., Kjeldsen, K.K., Tesakov, A.S., Snowball, I., Gosse, J.C., Alsos, I.G., Wang, Y., Dockter, C., Rasmussen, M., Jørgensen, M.E., Skadhauge, B., Prohaska, A., Kristensen, J.Å., Bjerager, M., Allentoft, M.E., Coissac, E., Rouillard, A., Simakova, A., Fernandez-Guerra, A., Bowler, C., Macias-Fauria, M., Vinner, L., Welch, J.J., Hidy, A.J., Sikora, M., Collins, M.J., Durbin, R., Larsen, N.K., Willerslev, E., 2022. A 2-million-year-old ecosystem in Greenland uncovered by environmental DNA. Nature 612, 283–291. 10.1038/s41586-022-05453-y

Langmead, B., Salzberg, S.L., 2012. Fast gapped-read alignment with Bowtie 2. Nat. Methods 9, 357–359. 10.1038/nmeth.1923

Lee, J., Kim, S., Jung, H., Koo, B.-K., Han, J.A., Lee, H.-S., 2023. Exploiting Bacterial Genera as Biocontrol Agents: Mechanisms, Interactions and Applications in Sustainable Agriculture. J. Plant Biol. 66, 485–498. 10.1007/s12374-023-09404-6

Lennon, J.T., Jones, S.E., 2011. Microbial seed banks: the ecological and evolutionary implications of dormancy. Nat. Rev. Microbiol. 9, 119–130. 10.1038/nrmicro2504

Li, D., Liu, C.-M., Luo, R., Sadakane, K., Lam, T.-W., 2015. MEGAHIT: an ultra-fast single-node solution for large and complex metagenomics assembly via succinct de Bruijn graph. Bioinformatics 31, 1674–1676. 10.1093/bioinformatics/btv033

Lu, J., Breitwieser, F.P., Thielen, P., Salzberg, S.L., 2017. Bracken: estimating species abundance in metagenomics data. PeerJ Comput. Sci. 3, e104. 10.7717/peerj-cs.104

Luczkiewicz, A., Kotlarska, E., Artichowicz, W., Tarasewicz, K., Fudala-Ksiazek, S., 2015. Antimicrobial resistance of Pseudomonas spp. isolated from wastewater and wastewater-impacted marine coastal zone. Environ. Sci. Pollut. Res. Int. 22, 19823–19834. 10.1007/s11356-015-5098-y

Lv, L., Wan, M., Wang, C., Gao, X., Yang, Q., Partridge, S.R., Wang, Y., Zong, Z., Doi, Y., Shen, J., Jia, P., Song, Q., Zhang, Q., Yang, J., Huang, X., Wang, M., Liu, J.-H., 2020. Emergence of a Plasmid-Encoded Resistance-Nodulation-Division Efflux Pump Conferring Resistance to Multiple Drugs, Including Tigecycline, in Klebsiella pneumoniae. mBio 11, e02930–19. 10.1128/mBio.02930-19

Mak, S., Xu, Y., Nodwell, J.R., 2014. The expression of antibiotic resistance genes in antibiotic-producing bacteria. Mol. Microbiol. 93, 391–402. 10.1111/mmi.12689

Martin, C., Timm, J., Rauzier, J., Gomez-Lus, R., Davies, J., Gicquel, B., 1990. Transposition of an antibiotic resistance element in mycobacteria. Nature 345, 739–743. 10.1038/345739a0

Martínez, J.L., 2008. Antibiotics and Antibiotic Resistance Genes in Natural Environments. Science 321, 365–367. 10.1126/science.1159483

Mikheenko, A., Prjibelski, A., Saveliev, V., Antipov, D., Gurevich, A., 2018. Versatile genome assembly evaluation with QUAST-LG. Bioinformatics 34, i142–i150. 10.1093/bioinformatics/bty266

Mohanty, S., Mishra, S., Saswat, S., Mohanty, B., 2026. Interconnected reservoirs and escalating resistance in multidrug-resistant Pseudomonas aeruginosa: An One Health review. Microb. Pathog. 215, 108478. 10.1016/j.micpath.2026.108478

Nolan, T.M., Reynolds, L.J., Sala-Comorera, L., Martin, N.A., Stephens, J.H., O’Hare, G.M.P., O’Sullivan, J.J., Meijer, W.G., 2023. Land use as a critical determinant of faecal and antimicrobial resistance gene pollution in riverine systems. Sci. Total Environ. 871, 162052. 10.1016/j.scitotenv.2023.162052

Olishevska, S., Nickzad, A., Déziel, E., 2019. Bacillus and Paenibacillus secreted polyketides and peptides involved in controlling human and plant pathogens. Appl. Microbiol. Biotechnol. 103, 1189–1215. 10.1007/s00253-018-9541-0

Parks, D.H., Chuvochina, M., Chaumeil, P.-A., Rinke, C., Mussig, A.J., Hugenholtz, P., 2020. A complete domain-to-species taxonomy for Bacteria and Archaea. Nat. Biotechnol. 38, 1079–1086. 10.1038/s41587-020-0501-8

Pärnänen, K., Karkman, A., Hultman, J., Lyra, C., Bengtsson-Palme, J., Larsson, D.G.J., Rautava, S., Isolauri, E., Salminen, S., Kumar, H., Satokari, R., Virta, M., 2018. Maternal gut and breast milk microbiota affect infant gut antibiotic resistome and mobile genetic elements. Nat. Commun. 9, 3891. 10.1038/s41467-018-06393-w

Pelegrino, K. de O., Campos, J.C., Sampaio, S.C.F., Lezirovitz, K., Seco, B.M., Pereira, M. de O., Rocha, D.A. da C., Jové, T., Nicodemo, A.C., Sampaio, J.L.M., 2015. fosI Is a New Integron-Associated Gene Cassette Encoding Reduced Susceptibility to Fosfomycin. Antimicrob. Agents Chemother. 60, 686–688. 10.1128/AAC.02437-15

Peterson, E., Kaur, P., 2018a. Antibiotic Resistance Mechanisms in Bacteria: Relationships Between Resistance Determinants of Antibiotic Producers, Environmental Bacteria, and Clinical Pathogens. Front. Microbiol. 9. 10.3389/fmicb.2018.02928

Peterson, E., Kaur, P., 2018b. Antibiotic Resistance Mechanisms in Bacteria: Relationships Between Resistance Determinants of Antibiotic Producers, Environmental Bacteria, and Clinical Pathogens. Front. Microbiol. 9. 10.3389/fmicb.2018.02928

Pochon, Z., Bergfeldt, N., Kırdök, E., Vicente, M., Naidoo, T., van der Valk, T., Altınışık, N.E., Krzewińska, M., Dalén, L., Götherström, A., Mirabello, C., Unneberg, P., Oskolkov, N., 2023. aMeta: an accurate and memory-efficient ancient metagenomic profiling workflow. Genome Biol. 24, 242. 10.1186/s13059-023-03083-9

Schlatter, D.C., Kinkel, L.L., 2014. Global biogeography of Streptomyces antibiotic inhibition, resistance, and resource use. FEMS Microbiol. Ecol. 88, 386–397. 10.1111/1574-6941.12307

Sim, S.B., Corpuz, R.L., Simmonds, T.J., Geib, S.M., 2022. HiFiAdapterFilt, a memory efficient read processing pipeline, prevents occurrence of adapter sequence in PacBio HiFi reads and their negative impacts on genome assembly. BMC Genomics 23, 157. 10.1186/s12864-022-08375-1

Simon, M.A., Ongpipattanakul, C., Nair, S.K., van der Donk, W.A., 2021. Biosynthesis of fosfomycin in pseudomonads reveals an unexpected enzymatic activity in the metallohydrolase superfamily. Proc. Natl. Acad. Sci. U. S. A. 118, e2019863118. 10.1073/pnas.2019863118

Surette, M.D., Wright, G.D., 2017. Lessons from the Environmental Antibiotic Resistome. Annu. Rev. Microbiol. 71, 309–329. 10.1146/annurev-micro-090816-093420

Tamminen, M., Virta, M., Fani, R., Fondi, M., 2012. Large-Scale Analysis of Plasmid Relationships through Gene-Sharing Networks. Mol. Biol. Evol. 29, 1225–1240. 10.1093/molbev/msr292

Uotila, K., Helamaa, M., Haggrén, G., Niemelä, T. (Eds.), 2025. Uuden Torin kantilla 1650–1827. Vol. 2: Turun Kauppatorin arkeologiset tutkimukset vuosina 2018–2022, 1. painos. ed, Kåkenhus-kirjat. Muuritutkimus Oy, Kaarina.

Viitamäki, S., Pessi, I.S., Virkkala, A.-M., Niittynen, P., Kemppinen, J., Eronen-Rasimus, E., Luoto, M., Hultman, J., 2022. The activity and functions of soil microbial communities in the Finnish sub-Arctic vary across vegetation types. FEMS Microbiol. Ecol. 98, fiac079. 10.1093/femsec/fiac079

Visuri, M., Nystrand, M., Auri, J., Österholm, P., Nilivaara, R., Boman, A., Räisänen, J., Mattbäck, S., Korhonen, A., Ihme, R., 2021. Maastokäyttöisten tunnistusmenetelmien kehittäminen happamille sulfaattimaille. Tunnistus-hankkeen loppuraportti.

Wang, C.-Z., Gao, X., Yang, Q.-W., Lv, L.-C., Wan, M., Yang, J., Cai, Z.-P., Liu, J.-H., 2021. A Novel Transferable Resistance-Nodulation-Division Pump Gene Cluster, tmexCD2-toprJ2, Confers Tigecycline Resistance in Raoultella ornithinolytica. Antimicrob. Agents Chemother. 65, e02229–20. 10.1128/AAC.02229-20

Wang, D., Zhan, Y., Cai, D., Li, X., Wang, Q., Chen, S., 2018. Regulation of the Synthesis and Secretion of the Iron Chelator Cyclodipeptide Pulcherriminic Acid in Bacillus licheniformis. Appl. Environ. Microbiol. 84, e00262–18. 10.1128/AEM.00262-18

Warinner, C., Herbig, A., Mann, A., Yates, J.A.F., Weiß, C.L., Burbano, H.A., Orlando, L., Krause, J., 2017. A Robust Framework for Microbial Archaeology. Annu. Rev. Genomics Hum. Genet. 18, 321–356. 10.1146/annurev-genom-091416-035526

Wellington, E.M., Boxall, A.B., Cross, P., Feil, E.J., Gaze, W.H., Hawkey, P.M., Johnson-Rollings, A.S., Jones, D.L., Lee, N.M., Otten, W., Thomas, C.M., Williams, A.P., 2013. The role of the natural environment in the emergence of antibiotic resistance in Gram-negative bacteria. Lancet Infect. Dis. 13, 155–165. 10.1016/S1473-3099(12)70317-1

Wibowo, M.C., Yang, Z., Borry, M., Hübner, A., Huang, K.D., Tierney, B.T., Zimmerman, S., Barajas-Olmos, F., Contreras-Cubas, C., García-Ortiz, H., Martínez-Hernández, A., Luber, J.M., Kirstahler, P., Blohm, T., Smiley, F.E., Arnold, R., Ballal, S.A., Pamp, S.J., Russ, J., Maixner, F., Rota-Stabelli, O., Segata, N., Reinhard, K., Orozco, L., Warinner, C., Snow, M., LeBlanc, S., Kostic, A.D., 2021a. Reconstruction of ancient microbial genomes from the human gut. Nature 594, 234–239. 10.1038/s41586-021-03532-0

Wibowo, M.C., Yang, Z., Borry, M., Hübner, A., Huang, K.D., Tierney, B.T., Zimmerman, S., Barajas-Olmos, F., Contreras-Cubas, C., García-Ortiz, H., Martínez-Hernández, A., Luber, J.M., Kirstahler, P., Blohm, T., Smiley, F.E., Arnold, R., Ballal, S.A., Pamp, S.J., Russ, J., Maixner, F., Rota-Stabelli, O., Segata, N., Reinhard, K., Orozco, L., Warinner, C., Snow, M., LeBlanc, S., Kostic, A.D., 2021b. Reconstruction of ancient microbial genomes from the human gut. Nature 594, 234–239. 10.1038/s41586-021-03532-0

Wood, D.E., Lu, J., Langmead, B., 2019. Improved metagenomic analysis with Kraken 2. Genome Biol. 20, 257. 10.1186/s13059-019-1891-0

Wright, G.D., 2010. Antibiotic resistance in the environment: a link to the clinic? Curr. Opin. Microbiol., Antimicrobials/Genomics 13, 589–594. 10.1016/j.mib.2010.08.005

Wright, M.S., Baker-Austin, C., Lindell, A.H., Stepanauskas, R., Stokes, H.W., McArthur, J.V., 2008. Influence of industrial contamination on mobile genetic elements: class 1 integron abundance and gene cassette structure in aquatic bacterial communities. ISME J. 2, 417–428. 10.1038/ismej.2008.8

Yakimov, M.M., Abraham, W.-R., Meyer, H., Laura Giuliano, Golyshin, P.N., 1999. Structural characterization of lichenysin A components by fast atom bombardment tandem mass spectrometry. Biochim. Biophys. Acta BBA - Mol. Cell Biol. Lipids 1438, 273–280. 10.1016/S1388-1981(99)00058-X

Zdouc, M.M., Blin, K., Louwen, N.L.L., Navarro, J., Loureiro, C., Bader, C.D., Bailey, C.B., Barra, L., Booth, T.J., Bozhüyük, K.A.J., Cediel-Becerra, J.D.D., Charlop-Powers, Z., Chevrette, M.G., Chooi, Y.H., D’Agostino, P.M., de Rond, T., Del Pup, E., Duncan, K.R., Gu, W., Hanif, N., Helfrich, E.J.N., Jenner, M., Katsuyama, Y., Korenskaia, A., Krug, D., Libis, V., Lund, G.A., Mantri, S., Morgan, K.D., Owen, C., Phan, C.-S., Philmus, B., Reitz, Z.L., Robinson, S.L., Singh, K.S., Teufel, R., Tong, Y., Tugizimana, F., Ulanova, D., Winter, J.M., Aguilar, C., Akiyama, D.Y., Al-Salihi, S.A.A., Alanjary, M., Alberti, F., Aleti, G., Alharthi, S.A., Rojo, M.Y.A., Arishi, A.A., Augustijn, H.E., Avalon, N.E., Avelar-Rivas, J.A., Axt, K.K., Barbieri, H.B., Barbosa, J.C.J., Barboza Segato, L.G., Barrett, S.E., Baunach, M., Beemelmanns, C., Beqaj, D., Berger, T., Bernaldo-Agüero, J., Bettenbühl, S.M., Bielinski, V.A., Biermann, F., Borges, R.M., Borriss, R., Breitenbach, M., Bretscher, K.M., Brigham, M.W., Buedenbender, L., Bulcock, B.W., Cano-Prieto, C., Capela, J., Carrion, V.J., Carter, R.S., Castelo-Branco, R., Castro-Falcón, G., Chagas, F.O., Charria-Girón, E., Chaudhri, A.A., Chaudhry, V., Choi, H., Choi, Y., Choupannejad, R., Chromy, J., Donahey, M.S.C., Collemare, J., Connolly, J.A., Creamer, K.E., Crüsemann, M., Cruz, A.A., Cumsille, A., Dallery, J.-F., Damas-Ramos, L.C., Damiani, T., de Kruijff, M., Martín, B.D., Sala, G.D., Dillen, J., Doering, D.T., Dommaraju, S.R., Durusu, S., Egbert, S., Ellerhorst, M., Faussurier, B., Fetter, A., Feuermann, M., Fewer, D.P., Foldi, J., Frediansyah, A., Garza, E.A., Gavriilidou, A., Gentile, A., Gerke, J., Gerstmans, H., Gomez-Escribano, J.P., González-Salazar, L.A., Grayson, N.E., Greco, C., Gomez, J.E.G., Guerra, S., Flores, S.G., Gurevich, A., Gutiérrez-García, K., Hart, L., Haslinger, K., He, B., Hebra, T., Hemmann, J.L., Hindra, H., Höing, L., Holland, D.C., Holme, J.E., Horch, T., Hrab, P., Hu, J., Huynh, T.-H., Hwang, J.-Y., Iacovelli, R., Iftime, D., Iorio, M., Jayachandran, S., Jeong, E., Jing, J., Jung, J.J., Kakumu, Y., Kalkreuter, E., Kang, K.B., Kang, S., Kim, W., Kim, G.J., Kim, H., Kim, H.U., Klapper, M., Koetsier, R.A., Kollten, C., Kovács, Á.T., Kriukova, Y., Kubach, N., Kunjapur, A.M., Kushnareva, A.K., Kust, A., Lamber, J., Larralde, M., Larsen, N.J., Launay, A.P., Le, N.-T.-H., Lebeer, S., Lee, B.T., Lee, K., Lev, K.L., Li, S.-M., Li, Y.-X., Licona-Cassani, C., Lien, A., Liu, J., Lopez, J.A.V., Machushynets, N.V., Macias, M.I., Mahmud, T., Maleckis, M., Martinez-Martinez, A.M., Mast, Y., Maximo, M.F., McBride, C.M., McLellan, R.M., Bhatt, K.M., Melkonian, C., Merrild, A., Metsä-Ketelä, M., Mitchell, D.A., Müller, A.V., Nguyen, G.-S., Nguyen, H.T., Niedermeyer, T.H.J., O’Hare, J.H., Ossowicki, A., Ostash, B.O., Otani, H., Padva, L., Paliyal, S., Pan, X., Panghal, M., Parade, D.S., Park, J., Parra, J., Rubio, M.P., Pham, H.T., Pidot, S.J., Piel, J., Pourmohsenin, B., Rakhmanov, M., Ramesh, S., Rasmussen, M.H., Rego, A., Reher, R., Rice, A.J., Rigolet, A., Romero-Otero, A., Rosas-Becerra, L.R., Rosiles, P.Y., Rutz, A., Ryu, B., Sahadeo, L.-A., Saldanha, M., Salvi, L., Sánchez-Carvajal, E., Santos-Medellin, C., Sbaraini, N., Schoellhorn, S.M., Schumm, C., Sehnal, L., Selem, N., Shah, A.D., Shishido, T.K., Sieber, S., Silviani, V., Singh, G., Singh, H., Sokolova, N., Sonnenschein, E.C., Sosio, M., Sowa, S.T., Steffen, K., Stegmann, E., Streiff, A.B., Strüder, A., Surup, F., Svenningsen, T., Sweeney, D., Szenei, J., Tagirdzhanov, A., Tan, B., Tarnowski, M.J., Terlouw, B.R., Rey, T., Thome, N.U., Torres Ortega, L.R., Tørring, T., Trindade, M., Truman, A.W., Tvilum, M., Udwary, D.W., Ulbricht, C., Vader, L., van Wezel, G.P., Walmsley, M., Warnasinghe, R., Weddeling, H.G., Weir, A.N.M., Williams, K., Williams, S.E., Witte, T.E., Rocca, S.M.W., Yamada, K., Yang, Dong, Yang, Dongsoo, Yu, J., Zhou, Z., Ziemert, N., Zimmer, L., Zimmermann, A., Zimmermann, C., van der Hooft, J.J.J., Linington, R.G., Weber, T., Medema, M.H., 2025. MIBiG 4.0: advancing biosynthetic gene cluster curation through global collaboration. Nucleic Acids Res. 53, D678–D690. 10.1093/nar/gkae1115

