## Supplemental_Material for "Antibiotic Resistomes And Microbial Communities In The 18th-Century Urban Settlement And Slaughterhouse Environment"

Supplementary Information

*Supplementary Table 1. Number of quality-trimmed classified and unclassified metagenomic reads after taxonomical classification with Kraken2*

| Sample | Classified_reads | Classified_% | Unclassified_reads | Unclassified_% |
|---|---|---|---|---|
| **M6419_a** | 28486334 | 26.09 | 80697198 | 73.91 |
| **M9342_a** | 8269696 | 19.21 | 34776660 | 80.79 |
| **M9352_a** | 20101162 | 25.39 | 59076802 | 74.61 |
| **M9378_a** | 31791533 | 34.03 | 61620938 | 65.97 |
| **M9409_a** | 23292767 | 30.43 | 53262915 | 69.57 |
| **M9413_1_a** | 2768215 | 25.34 | 8155321 | 74.66 |
| **M9413_2_a** | 7313102 | 27.40 | 19376910 | 72.60 |
| **M9413_3_a** | 4181613 | 31.72 | 9002855 | 68.28 |
| **M9416_a** | 19477048 | 33.82 | 38118689 | 66.18 |
| **M9436_a** | 24608684 | 29.93 | 57604435 | 70.07 |
| **M9438_a** | 26989756 | 28.44 | 67896502 | 71.56 |
| **M9441_a** | 43356873 | 48.74 | 45603138 | 51.26 |
| **NC1** | 365 | 32.07 | 773 | 67.93 |

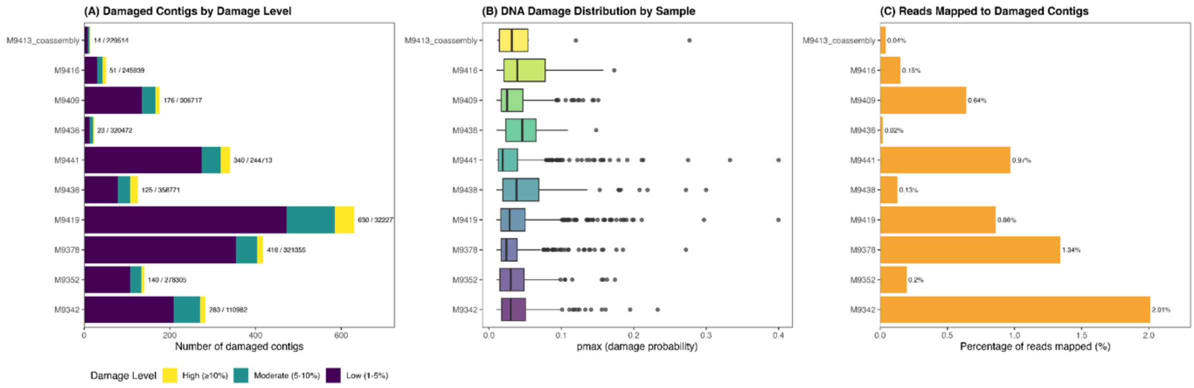


*Supplementary Figure 1. PyDamage analysis of ancient DNA damage patterns in metagenomic contigs. (A) Distribution of damaged contigs across samples colored by damage level. Bar height represents the total number of damaged contigs; stacked colors indicate damage categories based on pmax values: high damage (≥10%), moderate damage (5-10%), low damage (1-5%), and very low damage (<1%). Labels show the ratio of damaged to total contigs analyzed per sample (damaged/total). (B) DNA damage distribution by sample is shown as boxplots. Each point represents an individual damaged contig. The pmax value represents the modeled probability of C-to-T transitions at the 5' end of DNA fragments, characteristic of post-mortem*

*cytosine deamination. (C) Proportion of metagenomic reads mapped to damaged contigs, shown as percentages.*

*Supplementary Table 2. Mapped metagenomic reads against all contigs and damaged contigs extracted after pyDamage.*

| Reads against all contigs | | | | | | | |
|---|---|---|---|---|---|---|---|
| sample | total_reads | mapped_reads | percent_mapped | properly_paired | percent_properly_paired | singletons | percent_singletons |
| M6419_a | **218367064** | **177479225** | **81.28** | **167068472** | **paired** | **3028685** | **1.39** |
| M9342_a | **86092712** | **65918408** | **76.57** | **61543154** | **paired** | **1194008** | **1.39** |
| M9352_a | **158355928** | **110057358** | **69.50** | **103470406** | **paired** | **2801406** | **1.77** |
| M9378_a | **186824942** | **123513639** | **66.11** | **113781288** | **paired** | **3871567** | **2.07** |
| M9409_a | **153111364** | **104071623** | **67.97** | **95459344** | **paired** | **3674531** | **2.40** |
| M9413_1_a | **21847072** | **7453708** | **34.12** | **6555368** | **paired** | **615132** | **2.82** |
| M9413_2_a | **53380024** | **24401444** | **45.71** | **21523940** | **paired** | **1723156** | **3.23** |
| M9413_3_a | **26368936** | **9436719** | **35.79** | **8100792** | **paired** | **858571** | **3.26** |
| M9413_coassembly | **101596032** | **53608693** | **52.77** | **47847318** | **paired** | **3174509** | **3.12** |
| M9416_a | **115191474** | **75925675** | **65.91** | **69253778** | **paired** | **3023867** | **2.63** |
| M9436_a | **164426238** | **97273757** | **59.16** | **88220404** | **paired** | **4565533** | **2.78** |
| M9438_a | **189772516** | **135617203** | **71.46** | **126404026** | **paired** | **4202207** | **2.21** |
| M9441_a | **177920022** | **123169213** | **69.23** | **117899904** | **paired** | **3151423** | **1.77** |
| NC1 | **2276** | **9** | **0.40** | **2** | **paired** | **7** | **0.31** |

| | | | | | | | |
|---|---|---|---|---|---|---|---|
| Reads against damaged contigs | | | | | | | |
| M6419_a | **218367064** | **1885418** | **0.86** | **1604582** | **paired** | **237788** | **0.11** |
| M9342_a | **86092712** | **1728444** | **2.01** | **1471448** | **paired** | **210132** | **0.24** |
| M9352_a | **158355928** | **315435** | **0.20** | **262954** | **paired** | **46345** | **0.03** |
| M9378_a | **186824942** | **2506875** | **1.34** | **2168760** | **paired** | **261235** | **0.14** |
| M9409_a | **153111364** | **987183** | **0.64** | **823762** | **paired** | **137047** | **0.09** |
| M9413_1_a | **21847072** | **8129** | **0.04** | **6716** | **paired** | **1153** | **0.01** |
| M9413_2_a | **53380024** | **23338** | **0.04** | **18608** | **paired** | **3936** | **0.01** |
| M9413_3_a | **26368936** | **12262** | **0.05** | **8966** | **paired** | **2942** | **0.01** |
| M9413_coassembly | **101596032** | **43729** | **0.04** | **34290** | **paired** | **8031** | **0.01** |
| M9416_a | **115191474** | **170509** | **0.15** | **131540** | **paired** | **34119** | **0.03** |

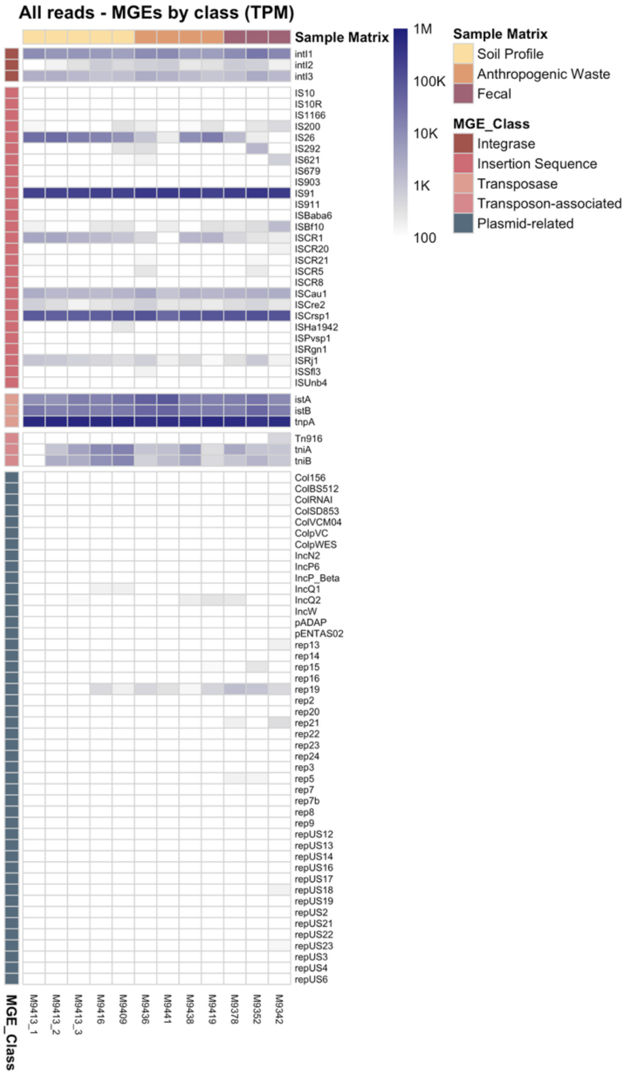


*Supplementary Figure 2. Heatmap showing all mobile genetic elements (MGEs) detected across 12 archaeological samples using BLASTN against MGE database (Pärnänen et al. 2018). Samples are displayed on the x-axis and detected gene families on the y-axis. BLASTN hits were retained at a percentage identity ≥ 80%, alignment length ≥ 40 bp, and bitscore ≥ 50. Read counts were normalized using Transcripts Per Million (TPM), and colour scales represent TPM values of 100–1,000,000. Prior to visualization, sequence variants of the same gene were grouped into gene families by removing the variant suffixes, and TPM values were calculated for the variants within each gene family. White cells indicate absence of detection.*

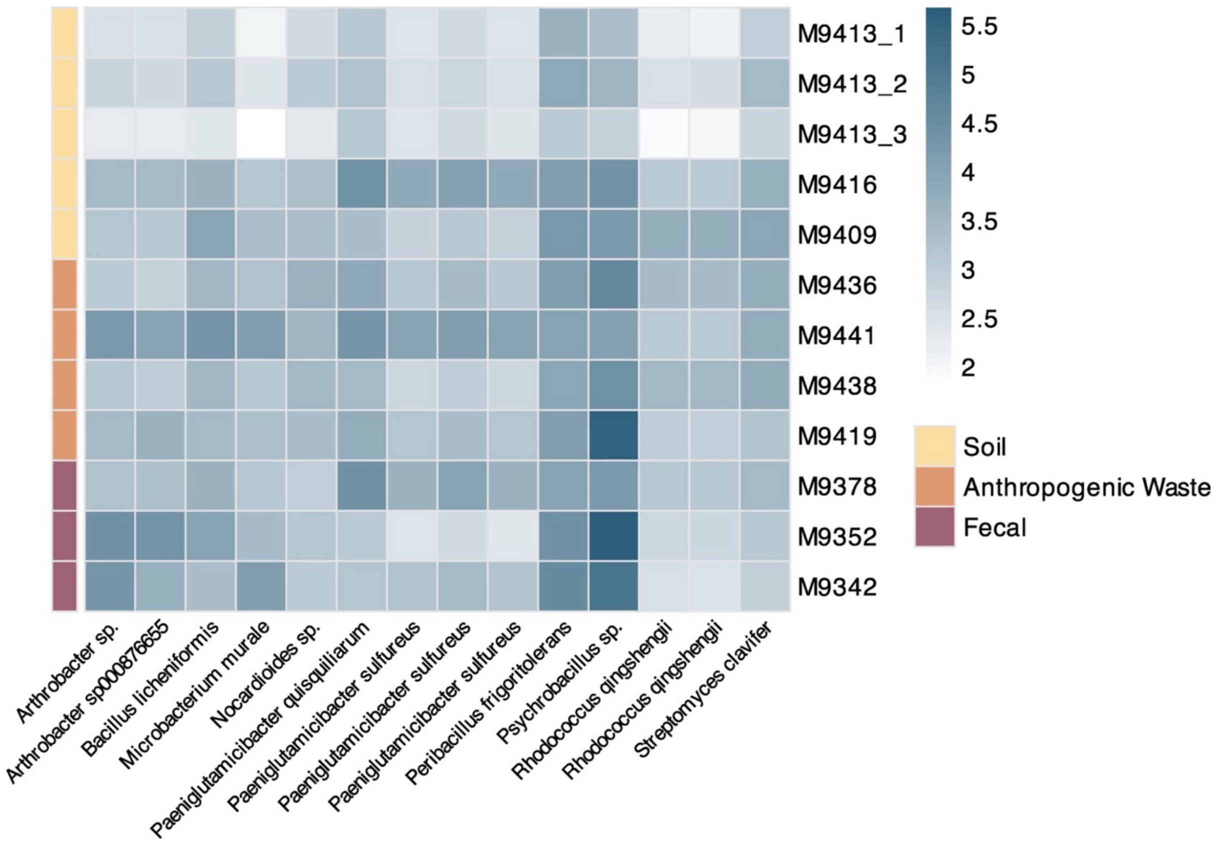


*Supplementary Figure 3. Heatmap of metagenomic read counts mapped to bacterial isolate genomes using Bowtie2. Rows represent the 12 metagenomic samples grouped by sample matrix (yellow: Soil; orange: Anthropogenic Waste; mauve: Fecal), and columns represent isolates cultivated from the 18th-century archaeological samples. The color scale represents $log_{10}$(count + 1)-transformed read counts, and white cells indicate no detected reads.*

*Supplementary Table 3. The genomic locations of detected ARGs and MGEs in isolate genomes.*

| isolate | contig | q_start | q_end | gene | gene_class | percent_identity | coverage | data_type | isolate_label |
|---|---|---|---|---|---|---|---|---|---|
| Arthrobacter sp. | **ptg000001l** | **780951** | **781576** | **ole(C)** | **Macrolide** | **70.827** | **65.5** | **ARG** | **Arthrobacter sp. (M9409)** |
| Nocardioides sp. | **ptg000001c** | **690246** | **690523** | **ARR-7** | **Rifamycin** | **72.598** | **62** | **ARG** | **Nocardioides sp. (M9419)** |
| Nocardioides sp. | **ptg000001c** | **2544419** | **2545684** | **tet(43)** | **Tetracycline** | **75.058** | **82.2** | **ARG** | **Nocardioides sp. (M9419)** |
| Bacillus licheniformis | **ptg000001c** | **3647882** | **3648447** | **blaZ** | **Beta-lactam** | **75.35** | **60.1** | **ARG** | **Bacillus licheniformis (M9378)** |
| Bacillus licheniformis | **ptg000001c** | **984773** | **985629** | **erm(D)** | **MLS** | **95.683** | **99.2** | **ARG** | **Bacillus licheniformis (M9378)** |
| Peribacillus frigoritolerans | **ptg000001c** | **5305415** | **5305983** | **blaZ** | **Beta-lactam** | **77.058** | **60** | **ARG** | **Peribacillus frigoritolerans (M9378)** |
| Peribacillus frigoritolerans | **ptg000001c** | **5364656** | **5365267** | **cat** | **Chloramphenicol** | **81.553** | **90** | **ARG** | **Peribacillus frigoritolerans (M9378)** |
| Streptomyces clavifer | **ptg000001l** | **3424892** | **3425991** | **cmlV** | **Chloramphenicol** | **79.982** | **84.6** | **ARG** | **Streptomyces clavifer (M9342)** |
| Streptomyces clavifer | **ptg000001l** | **5563529** | **5564443** | **ole(C)** | **Macrolide** | **79.528** | **95.4** | **ARG** | **Streptomyces clavifer (M9342)** |
| Arthrobacter sp. | **ptg000001l** | **2678146** | **2679113** | **tnpA** | **Transposase** | **82.051** | **100.4** | **MGE** | **Arthrobacter sp. (M9409)** |
| Microbacterium murale | **ptg000001c** | **3762556** | **3763091** | **ISCau1** | **Insertion Sequence** | **70.35** | **44.8** | **MGE** | **Microbacterium murale (M9419)** |
| Microbacterium murale | **ptg000001c** | **752558** | **753634** | **tnpA** | **Transposase** | **98.886** | **100** | **MGE** | **Microbacterium murale (M9419)** |
| Nocardioides sp. | **ptg000001c** | **5254660** | **5255294** | **ISCau1** | **Insertion Sequence** | **72.91** | **53.3** | **MGE** | **Nocardioides sp. (M9419)** |
| Paeniglutamicibacter quisquiliarum | **ptg000001c** | **3148475** | **3149551** | **tnpA** | **Transposase** | **100** | **100** | **MGE** | **Paeniglutamicibacter quisquiliarum (M9419)** |
| Paeniglutamicibacter sulfureus_1 | **ptg000002c** | **160036** | **161952** | **tnpA** | **Transposase** | **77.275** | **62.9** | **MGE** | **Paeniglutamicibacter sulfureus_1 (M9419)** |
| Paeniglutamicibacter sulfureus_2 | **ptg000001c** | **188720** | **190636** | **tnpA** | **Transposase** | **77.275** | **62.9** | **MGE** | **Paeniglutamicibacter sulfureus_2 (M9419)** |
| Paeniglutamicibacter sulfureus_3 | **ptg000002l** | **91470** | **93386** | **tnpA** | **Transposase** | **77.275** | **62.9** | **MGE** | **Paeniglutamicibacter sulfureus_3 (M9419)** |
| Rhodococcus qingshengii | **ptg000001c** | **6244196** | **6244731** | **ISCau1** | **Insertion Sequence** | **75.97** | **44.7** | **MGE** | **Rhodococcus qingshengii (M9352)** |
| Rhodococcus qingshengii | **ptg000002l** | **191091** | **191374** | **int2** | **Integrase** | **73.01** | **51.5** | **MGE** | **Rhodococcus qingshengii (M9352)** |

| | | | | | | | | | |
|---|---|---|---|---|---|---|---|---|---|
| Rhodococcus qingshengii | **ptg000003l** | **7685** | **7737** | **tniA** | **Transposon-associated protein** | **87.037** | **41.5** | **MGE** | **Rhodococcus qingshengii (M9352)** |
| Rhodococcus qingshengii | **ptg000003l** | **8666** | **9073** | **tniB** | **Transposon-associated protein** | **73.995** | **46.5** | **MGE** | **Rhodococcus qingshengii (M9352)** |
| Rhodococcus qingshengii | **ptg000004c** | **35634** | **38669** | **tnpA** | **Transposase** | **100** | **100** | **MGE** | **Rhodococcus qingshengii (M9352)** |
| Rhodococcus qingshengii_2 | **ptg000002c** | **1814207** | **1814742** | **ISCau1** | **Insertion Sequence** | **75.97** | **44.7** | **MGE** | **Rhodococcus qingshengii_2 (M9352)** |
| Rhodococcus qingshengii_2 | **ptg000002c** | **5387030** | **5387313** | **int2** | **Integrase** | **72.917** | **51.3** | **MGE** | **Rhodococcus qingshengii_2 (M9352)** |
| Streptomyces clavifer | **ptg000001l** | **3875823** | **3876457** | **ISCau1** | **Insertion Sequence** | **74.194** | **53.8** | **MGE** | **Streptomyces clavifer (M9342)** |
| Streptomyces clavifer | **ptg000002l** | **54946** | **55428** | **tnpA** | **Transposase** | **85.361** | **53** | **MGE** | **Streptomyces clavifer (M9342)** |
